# The PHD3-FOXO3 axis modulates the interferon type I response in microglia aggravating Alzheimer’s disease progression

**DOI:** 10.1101/2024.10.01.616066

**Authors:** Manuel A. Sanchez-Garcia, Nieves Lara-Ureña, Rosana March-Diaz, Clara Ortega-de San Luis, Silvia Quiñones-Cañete, Juan M. Barba-Reyes, Daniel Cabello-Rivera, Ana M. Muñoz-Cabello, Bella Mora-Romero, Carmen Romero-Molina, Antonio Heras-Garvin, Victoria Navarro, Jose Lopez-Barneo, Marisa Vizuete, Javier Vitorica, Ana B. Muñoz-Manchado, Matthew Cokman, Alicia E. Rosales-Nieves, Alberto Pascual

**Affiliations:** Instituto de Biomedicina de Sevilla (IBiS), Hospital Universitario Virgen del Rocio/CSIC/Universidad de Sevilla. 41013 Seville, Spain; Centro de Investigacion Biomedica en Red sobre Enfermedades Neurodegenerativas (CIBERNED), Madrid, Spain; Departamento de Neurociencias. Unidad de Biología Celular. Instituto de Investigación e Innovación Biomédica de Cádiz (INiBICA). University of Cádiz, Cádiz, Spain; Departamento de Fisiología Médica y Biofísica, Universidad de Sevilla, Seville, Spain; Departamento de Biología Celular, Facultad de Biología, Universidad de Sevilla, Seville, Spain; Departamento de Bioquímica y Biología Molecular, Facultad de Farmacia, Universidad de Sevilla; Hypoxia Biology Laboratory, Francis Crick Institute, London, NW1 1AT, UK

## Abstract

Microglia respond to Alzheimer’s disease (AD) with a variety of transcriptional responses. However, the regulation of specific transcriptional signatures and the contribution of each individual response to disease progression is only starting to be characterized. We have previously shown that hypoxia *via* hypoxia inducible factor 1 (HIF1) is a strong regulator of Aß plaque-associated microglia (AßAM). Here, we characterize the role of HIF1-mediated transcription of *Egln3*, encoding for PHD3, in AßAM. We show that oligomeric Aß treatment (oAß) *in vitro* induces the expression of *Hif1a* and *Egln3* in microglia, which correlates with the transcriptional activation of genes involved in the interferon type I signature (IFNS) in a PHD3-dependent manner. Mechanistically, we demonstrate FOXO3 to be an important repressor of IFNS in microglia, whose abundance decreases upon Aß presence, and, correspondingly, both in human single-nucleus (sn) and mouse AßAM transcriptomics, FOXO3 DNA binding sites define the IFNS. FOXO3 repression of the IFNS is dependent on PHD3, with our results suggesting a physical interaction between both proteins *in vitro*. *In vivo*, loss of PHD3 correlate with abrogation of the IFNS and activation of the disease-associated microglia signature (DAM) in AßAM. Transcriptional changes in microglia associate with increased microglia proximity to Aß plaques, augmented phagocytosis of Aß by microglia, reduced parenchymal levels of Aß, and an increase in small-sized plaques. PHD3 deficiency also reduced the Aß plaque-associated neuropathology and rescued behavioural deficits of an AD mouse model. Finally, we also demonstrate that microglial PHD3 overexpression during development in the absence of Aß pathology is sufficient to induce the IFNS and to behavioural alterations. Altogether, our data strongly indicate that the PHD3-FOXO3 axis controls the microglial IFNS in a cell autonomous manner, contributing to the progression of AD.

## Introduction

Microglia are brain-resident macrophages involved in several functions related with brain homeostasis, including myelin maintenance, efferocytosis of defective neurons, and vascular support^1,2^. Microglia are also involved in neurodegenerative disorders, contributing to the brain responses to local damage^2^. Analysis of genetic risk factors in Alzheimer’s disease (AD) has placed microglia in the spotlight of intense research by revealing that polymorphisms in genes with prominent or exclusive functions in microglia are linked to AD^3^. These polymorphisms are implicated in varied functional modules like lipid transport, endolysosomal functioning, and microglial migration and phagocytosis, among others^3^. In AD, microglia cluster^4^ and establish a protective barrier around senile plaques^5–8^, although they may also contribute to the progression of the disease by activating a proinflammatory phenotype^9–12^. As innate immune cells, microglia express a variety of cell-surface receptors^13^, which may bind to soluble amyloid ß peptide (Aβ) oligomers and fibrils^11^ or to other molecules released from injured cells. Aß activates diverse signalling pathways, including a common microglial neurodegenerative phenotype observed in disease-associated microglia (DAM)^14–17^, and, among others, a type-I interferon (IFN)^18–22^ and a HIF1-dependent signatures^23^. However, how these transcriptional signatures are regulated, how they interact with each other, and which are the functional consequences of their activation are only starting to be deciphered.

We have previously shown that loss of endothelial cells around Aß plaques^24^ induces a HIF1-mediated transcriptional program in Aß plaque-associated microglia (AßAM)^23^ and that HIF1 activity under hypoxia limits the microglial defensive potential against the disease^23^. HIF1 is transcriptionally regulated by different signalling pathways, including the mechanistic/mammalian target of rapamycin (mTOR) pathway^25^, which has been involved in the activation of AßAM^23,26^. HIF1 protein stability is also modulated in an oxygen-dependent manner as, in the presence of oxygen, HIF prolyl hydroxylases (PHD1 to 3) hydroxylate HIF1*α* subunit in specific prolines, which are recognized by the von Hippel Lindau (VHL) protein that ubiquitinates HIF1*α* and targets it for proteasomal degradation^27^. Under low oxygen levels, HIF1*α* accumulates, translocates to the nucleus, and activates a complex cell-type-dependent transcriptional program^27^. In monocyte-derived cells, HIF1 transcribes the TNF*α* and IL-1ß mRNAs, contributing to their proinflammatory activity^28,29^, and induces a switch from oxidation to fermentation of glucose^28^, a metabolic adaptation that has also been proposed in AßAM dysfunction^30,31^. Interestingly, a recent article has postulated HIF1, together with FOXO3 and FOXP2, as a main player in the transition between low to high inflammatory profile in human induced pluripotent stem cell (iPSC)-derived microglia^32^. However, we lack *in vivo* data of the contribution of these transcription factors in AßAM. *Egln3*, encoding for PHD3, is a HIF1-target gene with known functions in the regulation of innate immunity^33–37^. This gene has previously been found as one of the most unregulated transcripts in ABAM^23^ and regulates the stability of FOXO3^38,39^. This work aims to characterize the contribution of HIF1, PHD3, and FOXO3 to the functional and transcriptional responses of AßAM and their involvement in Alzheimer progression.

## Results

### HIF1 regulates *Egln3* in AßAM without altering cytokine production

We have previously shown that AßAM express the HIF1/Hypoxia induced module (HMM) and that overactivation of HIF1 leads to microglial quiescence^23^. However, the role of HIF1 in homeostatic or activated microglia has not been addressed. In monocyte-derived cells, HIF1 is associated with the transcription of the proinflammatory cytokines IL1ß and TNF*α*^28,29^ and a similar role has been suggested in microglia^30,31,40^. To test the role of HIF1 in microglia, we generated a HIF1*α*-deficient mouse model with and without Aß deposition (*Cx3cr1^Cre::ERT2/+^*; *Hif1a^Flox/Flox^* ± *APP-PSEN1/+*). We treated 2-month-old mice with tamoxifen in the diet for 30 days, aged the mice to 12-month-old, isolated the microglia by fluorescence-activated cell sorting (FACS), and checked that *Hif1a* mRNA levels were strongly downregulated in both wildtype and Aß-depositing mouse models in tamoxifen-treated *Cx3cr1^Cre::ERT2/+^*; *Hif1a^Flox/Flox^* mice (Fig. 1a), a finding in line with previous observations in *Cx3cr1^Cre::ERT2/+^*; *Hif1a^Flox/Flox^* mouse primary microglial cultures^23^. Accordingly, *Hif1a* itself and genes belonging to the HMM were induced in AßAM and downregulated in HIF1 deficient microglia, including *Egln3*, *Vegfa*, *Bhlhe40*, *Ero1l*, and *Mxi1* (Fig. 1b). Surprisingly, *Tnf* was not induced by the pathology nor regulated by HIF1 and *Il1ß* was induced in AßAM but not regulated by HIF1 (Fig. 1c), strongly suggesting a different role of HIF1 in microglia.

**Figure 1.**
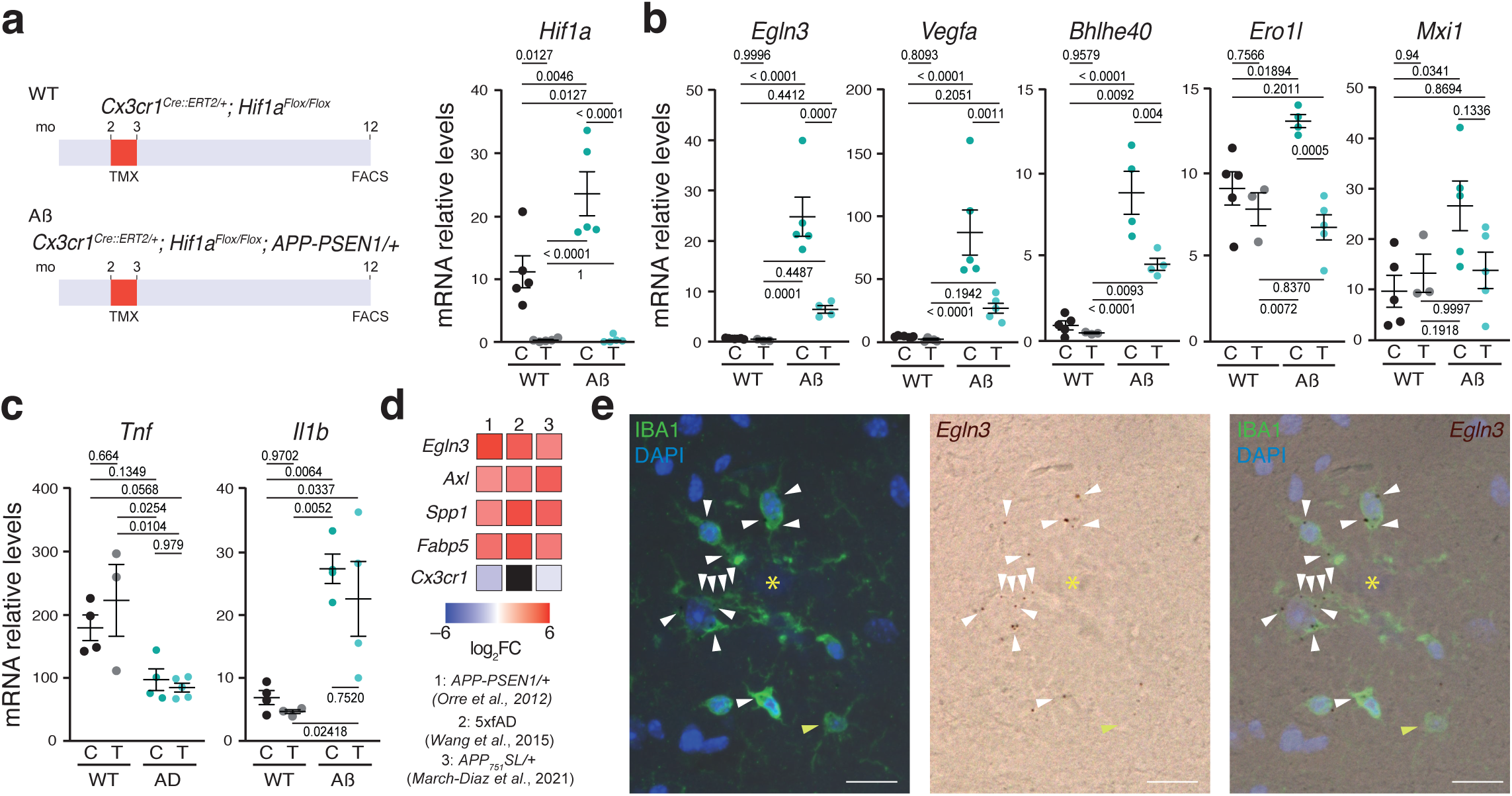
HIF1-mediated expression of *Egln3* in AßAM. **a**–**c**, Left panel, mouse models used. mo: -month-old. Right graphs, *Hif1a* (**a**) and HMM-included genes (**b**,**c**) mRNA expression in FACS-isolated microglia from 12-month-old control –C– and tamoxifen –T– treated wild-type (WT) and *APP-PSEN1/+* (Aß) mice was analysed by qRT-PCR. **d**, Heatmap of the log_2_ fold change (FC) of *Egln3*, *Axl*, *Spp1*, *Fabp5*, and *Cx3cr1* genes in three different Aß-depositing models. **e**, *In situ* hybridization of *Egln3* mRNA, immunohistochemistry for microglia (IBA1), and nuclear (DAPI) staining in brain sections of 12-month-old *APP-PSEN1/+* mice proximal (white arrowheads) and distal (yellow arrowheads) to Aβ plaques (yellow asterisk). Scale bars, 10 µm. All data are presented as means ± s.e.m. Individual points in the graphs indicate independent mice used per experiment. *p*-values from two-sided Student’s *t*-test.

To further investigate the role of HIF1 in AßAM, we focus on *Egln3*, a well-known HIF1-target gene^41^ that is involved in the modulation of HIF1 and FOXO3 stability^38,39,42^. In the immune system, PHD3 has a predominant role in the control of inflammation in several cell types including macrophages^33–37^, suggesting a role on HIF1/FOXO3-mediated control of innate immunity.

To further characterize the expression of PDH3 in microglia, we reanalysed the expression of *Egln3* mRNA in microglia isolated by FACS from several Aß-depositing models^16,23,43^ and compared with the expression changes in some genes highly expressed by AßAM (Fig. 1d). This approach enabled us to observe a substantial induction of *Egln3* across different mouse models. To confirm the transcriptomic data, we performed *in situ* hybridization combined with immunofluorescence. *Egln3* mRNA colocalized with IBA1 immunoreactive cells in the proximity of Aß deposits but showed low expression in distal-to-plaque homeostatic microglia (Fig. 1e), similar to what was observed in global transcriptomic analysis (Fig. 1d). Together, our data indicate that HIF1 contributes to the expression of PDH3 in AßAM.

### PHD3 regulates FOXO3, a repressor of the IFN microglia

As described before, PHD3 has a role in the innate immune system^33–37^, is strongly upregulated in AßAM, and interacts with and modifies the stability of FOXO3^38,39^, a protein that could be involved in the transition from low to high inflammatory microglia^32^. Mechanistically, FOXO3 represses the transcription of antiviral IFN target genes in macrophages through direct binding to their promoters^44^. IFN microglia is present in AD and have recently been characterized in models of AD and in humans^18–22^. Importantly, the activity of IFN stimulated genes has been associated with decreased synaptic markers and worsening of behavioural function in AD mouse models^20,45^ and with physiologic pruning of synapsis during development^46^. We therefore hypothesized that a HIF1-PHD3-FOXO3 axis may be controlling the IFNS expression in microglia (Fig. 2a). To understand the role of PHD3 in microglia, we exposed primary neonatal microglial cell cultures to oligomeric Aß (oAß). HIF1*α* was strongly induced both at mRNA and protein levels (Fig. 2b) and was accompanied by the augmentation of its target gene *Egln3* with a similar temporal profile (Fig. 2c). Then, we tested whether oAß treatment induced *in vitro* the expression of the IFNS, including the master regulator *Irf7*^47^, observing an induction with a similar kinetic to HIF1/PHD3 expression (Fig. 2d), which reproduce the induction of the IFNS observed in FACS-isolated AßAM from *5xfAD/+*, *APP-PSEN1/+*, *APP_751_SL/+*, and mouse models (Supplementary Fig. 1a). To evaluate the role of PHD3 in the upregulation of the IFNS by oAß, we performed primary neonatal cultures of control and PHD3-deficient microglia (*Egln3^−/−^* mice) and observed a significant downregulation of the IFNS in the absence of PHD3 (Fig. 2e). To further understand how PHD3 regulates the IFNS, we investigated whether oAß could modulate FOXO3 levels. Interestingly, oAß treatment reduced FOXO3 protein levels in wildtype (Fig. 3a) but not in *Egln3^−/−^* (Fig. 3b) microglia, strongly suggesting that PHD3 controls FOXO3 abundance. To corroborate that the levels of FOXO3 are directly linked with the expression of the IFNS, we knockdown FOXO3 expression in primary neonatal microglial cell cultures. Knockdown of FOXO3 was verified at the mRNA and protein levels (Fig. 3c,d) and resulted in the activation of the IFNS (Fig. 3e), suggesting that, like in peripheral macrophages^44^, FOXO3 represses IFNS transcription in microglia.

**Figure 2.**
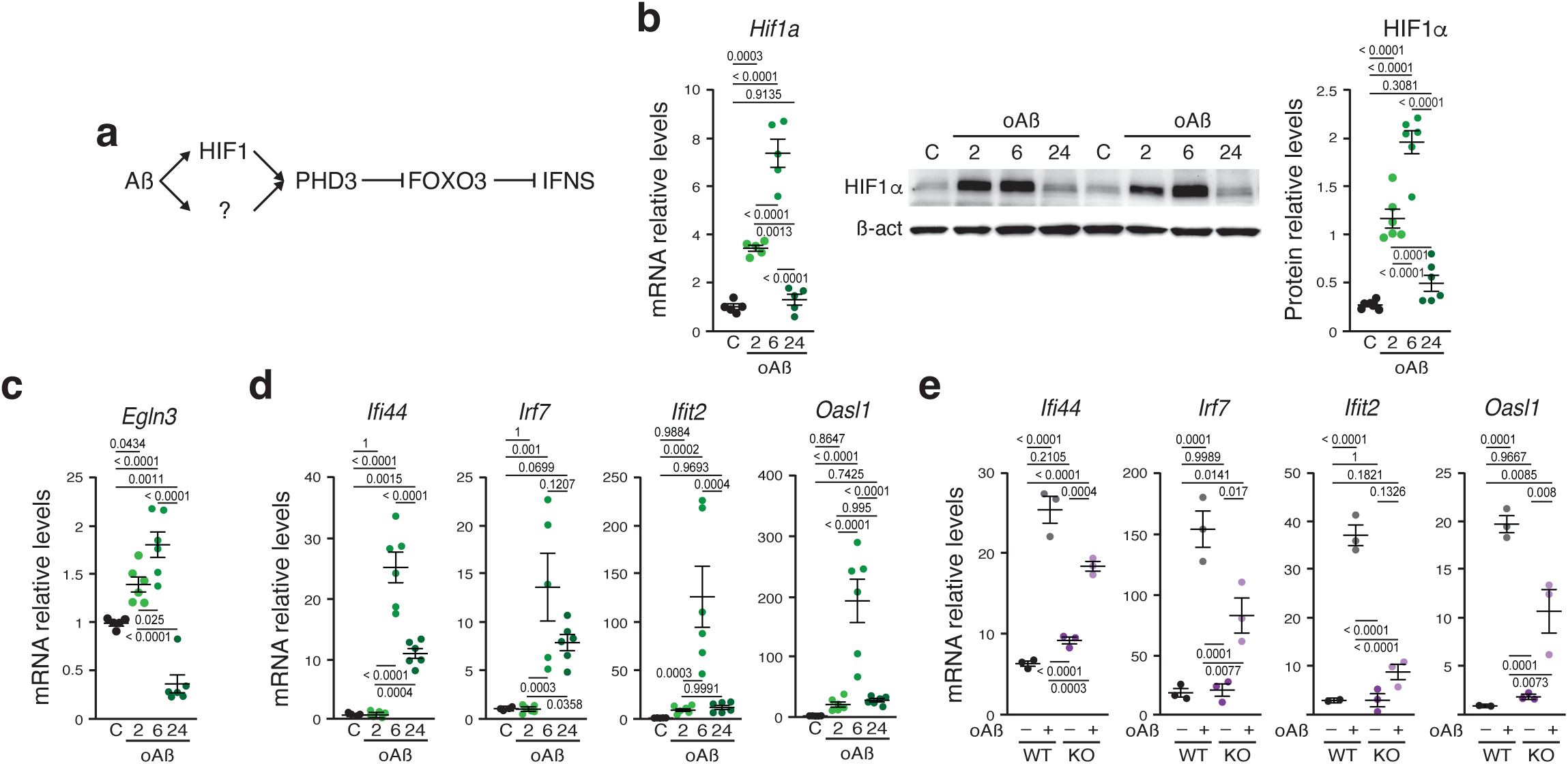
PHD3 regulates the microglial IFNS in response to Aß. **a**, Hypothetical pathway involved in the activation of IFNS by oAß. Arrows indicate activation, cross-lines indicate repression. **b**–**d**, *Hif1a* (**b**), *Egln3* (**c**), and IFNS (**d**) mRNAs; and HIF1*α*protein (**b**) levels in primary neonatal microglial cell cultures exposed to control (C) or oligomeric Aß (oAß) for 2, 6, or 24 h. *p*-values from ANOVA with Tukey’s post-test. **e**, mRNA levels of antiviral genes were estimated in primary neonatal microglial cell cultures from wild-type (WT) or *Egln3^−/−^* (KO) mice exposed to control (–) or oligomeric Aß (+). *p*-values from ANOVA with Tukey’s post-test.

**Figure 3.**
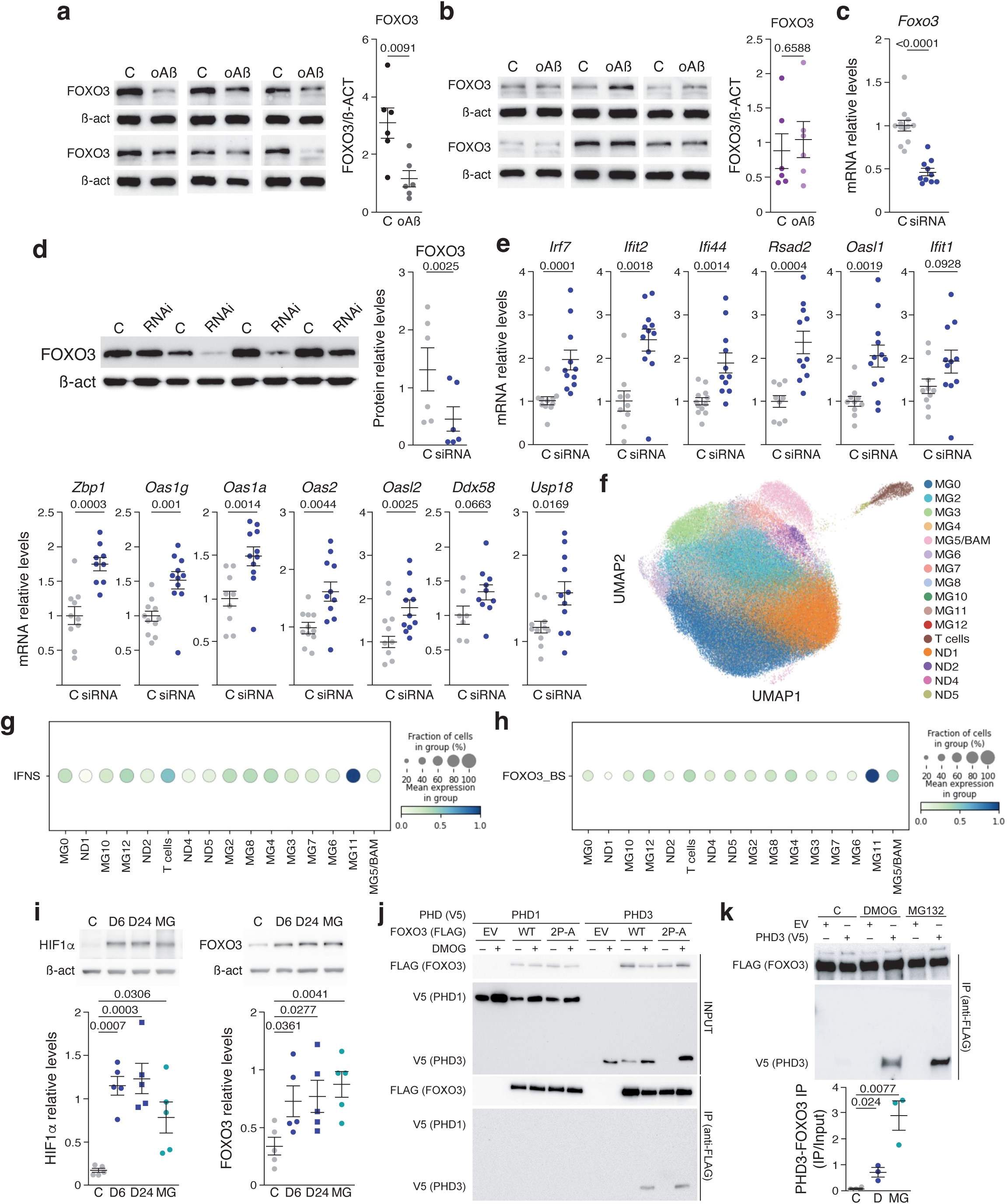
PHD3 regulates through FOXO3 the microglial IFNS. **a**,**b**, FOXO3 protein levels in primary neonatal microglial cell cultures from WT (**a**) and *Egln3^−/−^* (**b**) mice exposed to control (C) or oligomeric Aß (oAß). *p*-values from Student’s *t*-test. **c**,**d**, *Foxo3* mRNA (**c**) and FOXO3 protein (**d**) levels were estimated in primary microglial cell cultures exposed to control (C) or an anti-*Foxo3* siRNA. *p*-values from Mann-Whitney’s test (**a**) or two-sided Student’s *t*-test (**d**). **e**, IFNS mRNA levels in primary microglial cell cultures exposed as in (**a**,**b**). *p*-values from Mann-Whitney’s test for *Irf7*, *Ifit2, Oas1g*, and *Ddx58*; and two-sided Student’s *t*-test for *Ifi44*, *Rsad2*, *Oasl1*, *Ifit1*, *Zbp1*, *Oas1a, Oas2*, *Oasl2*, and *Usp18*. **f**, UMAP plot illustrating 11 distinct microglial states as identified by^32^. Each dot corresponds to an individual cell, sampled from six different brain regions: prefrontal cortex (PFC), hippocampus, mid-temporal cortex, angular gyrus, entorhinal cortex, and thalamus. Cells are clustered based on their transcriptional profiles, highlighting the heterogeneity of microglial states across these regions. **g,h**, Dot plot showing the expression of the IFN (**g**) and FOXO3 binding sites (BS) (**h**) gene signatures across various microglial states from the entorhinal cortex and hippocampus. Each circle represents the mean expression level (indicated by colour intensity) and the proportion of cells expressing the IFN or FOXO3_BS signatures within each state (reflected by dot size). Notably, MG11 displays a significant overexpression of both IFN (_log_FoldChange = 1.7; *p*-value < 0.05) and FOXO3_BS (_log_FoldChange = 2.03; *p*-value < 0.05) signatures. **i**, HIF1*α* (left) and FOXO3 (right) protein levels in primary neonatal microglial cell cultures exposed to control (C), DMOG (D; 6 or 24 h), or MG132 (MG; 24 h). *p*-values from ANOVA with Tukey’s post-test. **j**, HEK-293T cells transfected with both PHD1-V5 or PHD3-V5 and an empty (EV), wildtype (WT) FOXO3-FLAG, or P_426_A/P_437_A mutant (2P-A) FOXO3-FLAG vector; and incubated with (+) or without (-) DMOG for 24 h. Upper panels show western blots with anti-V5 (PHD1 and PHD3) and anti-FLAG (FOXO3) in proteins extracts (INPUT) and, bottom panels, after immunoprecipitation (IP) with an anti-FLAG antibody. **k**, HEK-293T cells transfected with both FOXO3-FLAG and an empty (EV) or a PHD3-V5 vector; and incubated with DMOG (+) or MG132 (+) for 24 h. Upper panels show western blots with anti-FLAG (FOXO3) and anti-V5 (PHD3) after immunoprecipitation (IP) with an anti-FLAG antibody. Bottom graph, quantification of upper panels. *p*-values from Student’s *t*-test. All data are presented as means ± s.e.m. Individual points in the graphs represent the *n* and are independent microglial primary (**a**–**e**) or HEK-293T (**i**,**k**) cultures.

To corroborate the role of FOXO3 regulating the transcription of the IFN microglia *in vivo*, we first reanalysed the transcriptomic data from acutely isolated AßAM from an AD mouse model and observed a prominent enrichment of the genes containing experimentally demonstrated FOXO3 DNA binding sites^44^ (Supplementary Fig. 1b). To validate the FOXO3 role in controlling gene expression of human IFN microglia, we reanalysed the single-cell (sc) RNAseq data from a previous study^32^. We clustered the cells using the same markers than in^32^ (Fig 3f) and clearly identified the IFN microglia (Fig. 3g; MG11). We then interrogated the expression of the genes controlled by FOXO3 in the different microglial types, observing a clear contribution of FOXO3 regulated genes to human IFN microglia (Fig. 3h).

PHDs may regulate FOXO3 stability through direct or indirect hydroxylation or by direct interaction^38,39^, therefore we evaluated if PHD activity could be controlling the levels of FOXO3 in microglia. To this end, we first exposed primary cultures of neonatal microglia to the pan-PHDs inhibitor DMOG and evaluated the levels of HIF1*α*, as a control, and FOXO3. FOXO3 was accumulated with a similar kinetic than HIF1*α* (Fig. 3i) suggesting that PHDs control directly or indirectly the levels of FOXO3 in microglia. Since FOXO3 protein levels are controlled by proteasomal degradation^38^, MG132 proteasome inhibitor was used to identify accumulation of HIF1*α* and FOXO3 in microglia (Fig. 3i). Finally, to evaluate whether a physical interaction exists between PHD3 and FOXO3 and its dependence on PHDs hydroxylase activity, we overexpressed both tagged-proteins in HEK-293T cells and included PHD1 as a control and performed co-immunoprecipitation studies. First, we validated the technique by using HIF1*α* and observed co-immunoprecipitation between both PHD1 and PHD3 and HIF1*α* after PHDs inhibition with DMOG (Supplementary Fig. 1c). Interestingly, FOXO3 interacted with PHD3 and not with PHD1 in the presence of DMOG (Fig. 3j,k and Supplementary Fig. 1e) or MG132 (Fig. 3j and Supplementary Fig. 1e; only PHD3), suggesting a specific interaction between both proteins. FOXO3 direct hydroxylation has not been reproduced *in vitro*^48^ but we observed a clear interaction with FOXO3 by inhibiting PHDs activity. To test whether the interaction was dependent of direct hydroxylation of prolines in FOXO3, we mutated the two prolines described as hydroxylated in FOXO3^38^ and observed the same interaction in the presence but not in the absence of DMOG (Fig. 3k), strongly suggesting that DMOG may be inhibiting the activity of PHDs over other targets or, alternatively, that DMOG treatment make any conformational change in PHD3 that stabilizes the interaction with FOXO3. Altogether, these results indicate that oAß induces HIF1 and PHD3, which reduces the stability of FOXO3 in primary microglial cultures leading to the upregulation of the IFNS.

### PHD3 deficiency abolishes the IFN response in AßAM

PHD3 has low expression in the brain^49^ and PHD3 inactivation has a minor systemic phenotype^50^, therefore, and to further understand the role of PHD3 in AßAM, we generated a mouse model combining lack of PHD3 with Aß deposition (*Egln3^−/−^*; *APP-PSEN1/+*). First, we compared the percentage of CD11b^+^/CD45^+^ microglia from 12-month-old wild-type, *Egln3^−/−^*, *APP-PSEN1/+*, and *Egln3^−/−^*; *APP-PSEN1/+* mice using flow cytometry (Supplementary Fig. 2a). No differences were observed between wild-type and *Egln3^−/−^* mice in the absence of Aß accumulation (Fig. 4a), in agreement with the low levels of PHD3 reported in non-reactive microglia (Fig. 1d). By contrast, the deposition of Aß in *Egln3^−/−^*; *APP-PSEN1/+* and *APP-PSEN1/+* mice induced a similar increase in the percentage of CD11b^+^/CD45^+^ microglial cells (Fig. 4a). This finding was further confirmed by stereology-assisted counting of IBA1-immunoreactive microglia in the cortex of control and PHD3-decificient Aß-depositing models (Fig. 4b). However, although the total number of microglial cells did not differ between *Egln3^−/−^*; *APP-PSEN1/+* and *APP-PSEN1/+* mice, microglial cells expressed higher levels of CD45 in AD mice lacking PHD3 (Fig. 4c), suggesting a change in activation status^51^.

Secondly, to characterize the activation state of PHD3-deficient microglia in an AD mouse model, we isolated microglia using FACS and performed global transcriptional expression profiling (Supplementary Table 1). As expected, *Egln3* was the most downregulated gene (Supplementary Table 1), and gene set enrichment analysis (GSEA) showed upregulation of the DAM signature (Fig. 4d and Supplementary Table 2). Correspondingly, genes strongly upregulated in AD mouse models^23^, including *Mamdc2*, *Postn*, and *Cxcl14*, were further increased by PHD3 deficiency(Fig. 4d), suggesting a different microglial activation.

**Figure 4.**
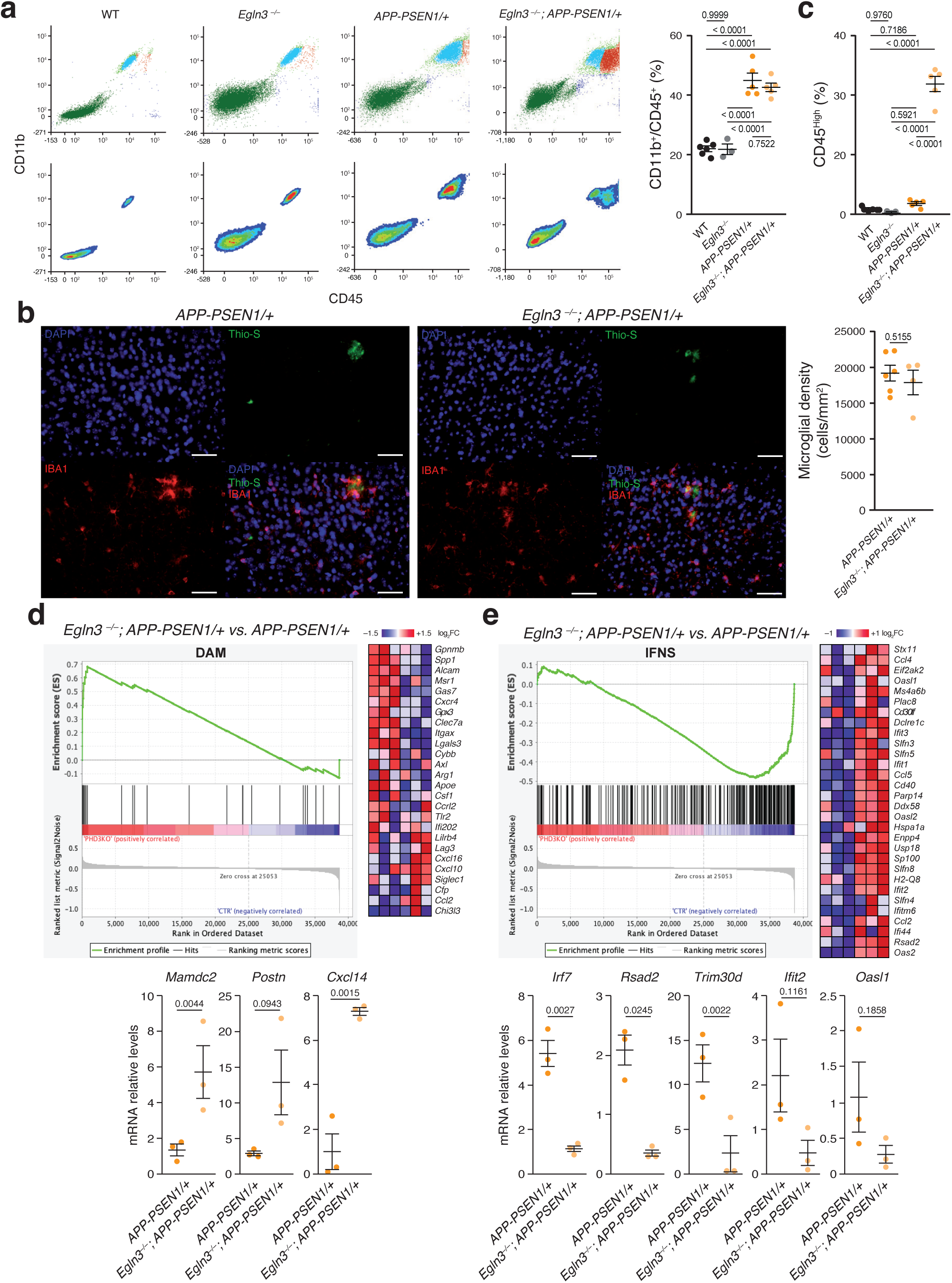
PHD3 regulates the IFNS in AßAM. **a**, Adult microglia from 12-month-old wild-type (WT), *Egln3^−/−^*, *APP-PSEN1/+* and *Egln3^−/−^*; *APP-PSEN1/+* mice were characterized by surface expression of CD45 and CD11b. In the upper panels, each dot represents one fluorescent event. Lower panels depict the density plots from the data in the upper panels. Right graph, quantification of the percentage of CD45/CD11b positive events. *p*-values from ANOVA with Tukey’s post-test. **b**, Stereological estimation of microglial cell number (#). Left, representative cortical coronal sections from *APP-PSEN1/+* and *Egln3^−/−^*; *APP-PSEN1/+* mice immunostained with an anti-IBA1 antibody and stained with Thioflavin-S (Thio-S). DAPI was used to label nuclei. Scale bars, 50µm. Right graph, number of IBA1 positive cells per hemicortex of *APP-PSEN1/+* and *Egln3^−/−^*; *APP-PSEN1/+* mice. *p*-values from two-sided Student’s *t*-test. **c**, Quantification of the percentage of CD45^High^ positive events in (**a**). Low and high populations were determined using the density plots shown in (**a**, lower panels). *p*-values from ANOVA with Tukey’s post-test. **d**,**e**, Upper panels, GSEA of FACS-isolated microglia from 12-month-old *Egln3^−/−^*; *APP-PSEN1/+ versus APP-PSEN1/+* showing DAM (**d**) and IFNS (**e**) signatures plots (left) and the heatmaps (right) of the top 30 ranking leading edge genes. FC: fold change. Bottom graphs, qRT-PCR analysis of the expression of several genes induced (left) or repressed (right) in *Egln3^−/−^*; *APP-PSEN1/+* microglia. *p*-values from Student’s *t*-test. All data are presented as means ± s.e.m. Individual points in the graphs indicate independent mice used per experiment.

To further understand the changes taking place in the microglia of PHD3-deficient AD mice, we focused on downregulated pathways. Microglia is characterized by a strong induction of several transcriptomic modules when responding to external or internal stimuli like neurodegeneration^23,52^. However, transcriptional downregulation is weakly associated with activation of microglia, with the exception of a few homeostatic genes^14,15,17^. According with our *in vitro* results, the most prominent gene sets (GSs) downregulated in *Egln3^−/−^*; *APP-PSEN1/+* mice included the Nod-like, Toll-like, and Rig-I-like receptor signalling pathways, and the cytosolic DNA sensing pathway. These pathways are related to antiviral responses mediated by IFN, which are upregulated in AßAM (Supplementary Fig. 2b and Supplementary Table 2). To statistically evaluate the relevance of the IFNS in PHD3 deficient AD microglia, we obtained an IFN-ß-induced signature in macrophages from a previous report^44^. GSEA revealed a strong downregulation of the IFN-ßS in AD mouse models lacking PHD3 (Fig. 3e and Supplementary Table 2). Finally, we observed a negative enrichment in the genes containing experimentally demonstrated FOXO3 DNA binding sites^44^ (Supplementary Fig. 2c), reinforcing the idea that FOXO3 represses IFNS *via* PHD3. We confirmed our microarray data using qRT-PCR (Fig. 4e) for a set of genes including *Irf7*, a key player in the establishment of the antiviral response^47^. Altogether, our data strongly suggest that PHD3 contributes to the induction of the IFNS in AßAM.

### PHD3 deficiency improves functional microglial responses to Aß

With the exception of the genes involved in the IFNS, the majority of genes regulated in the absence of PHD3 were genes already induced in AßAM, suggesting that PHD3 mutation increases the microglial responses to Aß. We speculate that the reduction of the antiviral module by PHD3 deficiency enables the up-regulation of other functional modules. To investigate the functional consequences of PHD3 deficiency in AßAM, we first evaluated two genes involved in the capacity of microglia to respond to Aß accumulation and linked genetically to AD, *Trem2*^53,54^ and *Cd33*^55^. A loss-of-function mutation in *Trem2* (R47H) is one of the strongest risk factors for AD and TREM2 deficiency associates with a worsening of AD models (for a review see^56^). Interestingly, PHD3 limits *Trem2* expression in AD microglia, as its absence produces a further induction of the gene (Fig. 5a). Conversely, PHD3 absence correlated with a decrease in the *Cd33* expression in microglia, whose elevated abundance in AD brains is known to limit microglial phagocytic capacity^55^, (Fig. 5b). Interestingly, full activation of the DAM signature depends on TREM2 activity^15^, a signature that contains genes related to the ability of microglia to cluster around and phagocyte Aß plaques^15,43^. We therefore tested if the microglial spatial disposition could be altered by PHD3 deficiency by calculating a proximity index to Aß plaques. We observed a significant increase in the microglial proximity index *Egln3^−/−^*; *APP-PSEN1/+* compared with *APP-PSEN1/+* mice (Fig. 5c). However, the number of microglial cells surrounding the Aß plaques (Supplementary Fig. 3a), the total microglial load (Supplementary Fig. 3b), or the morphology of microglia distal to Aß plaques (Supplementary Fig. 3c) was not altered in PHD3 deficient mice. To test whether phagocytosis of Aß could be altered by PHD3 deficiency, we measured the microglia uptake of Aß using *in vivo* labelling with methoxy-X04, a fluorescent compound that can cross the blood-brain barrier and possesses high binding affinity for Aß^10^. Flow cytometry analysis showed a trend to increase the proportion of phagocytic microglia in *Egln3^−/−^*; *APP-PSEN1/+* compared with *APP-PSEN1/+* mice (Fig. 5d). Correspondingly, a decrease was observed in the total cortical levels of monomeric Aß_1-42_ and a trend in Aß_1-40_ (Fig. 5e) by ELISA. To study Aß plaques deposition, we stained cortical brain sections with Thioflavin-S (Thio-S), which labels ß-sheet structure-containing proteins like Aß, and with an anti-Aß antibody. We first estimated the percentage of the cortex covered (Thio-S or Aß load) by dense-core Aß plaques observing no differences between groups (Fig. 5f and Supplementary Fig. 3d). However, we observed a trend to increase the total number of Aß plaques in PHD3 deficient mice but with a reduced average Aß plaque size (Fig. 5f). Size distribution revealed that PHD3 mice were characterized by an elevated abundance of small-sized Aß plaques with no changes in the bigger plaques (Fig. 5g). Interestingly, the uplift observed in Aß plaques burden could be attributed to core-dense and not filamentous Thio-S positive plaques (Fig. 5h).

**Figure 5.**
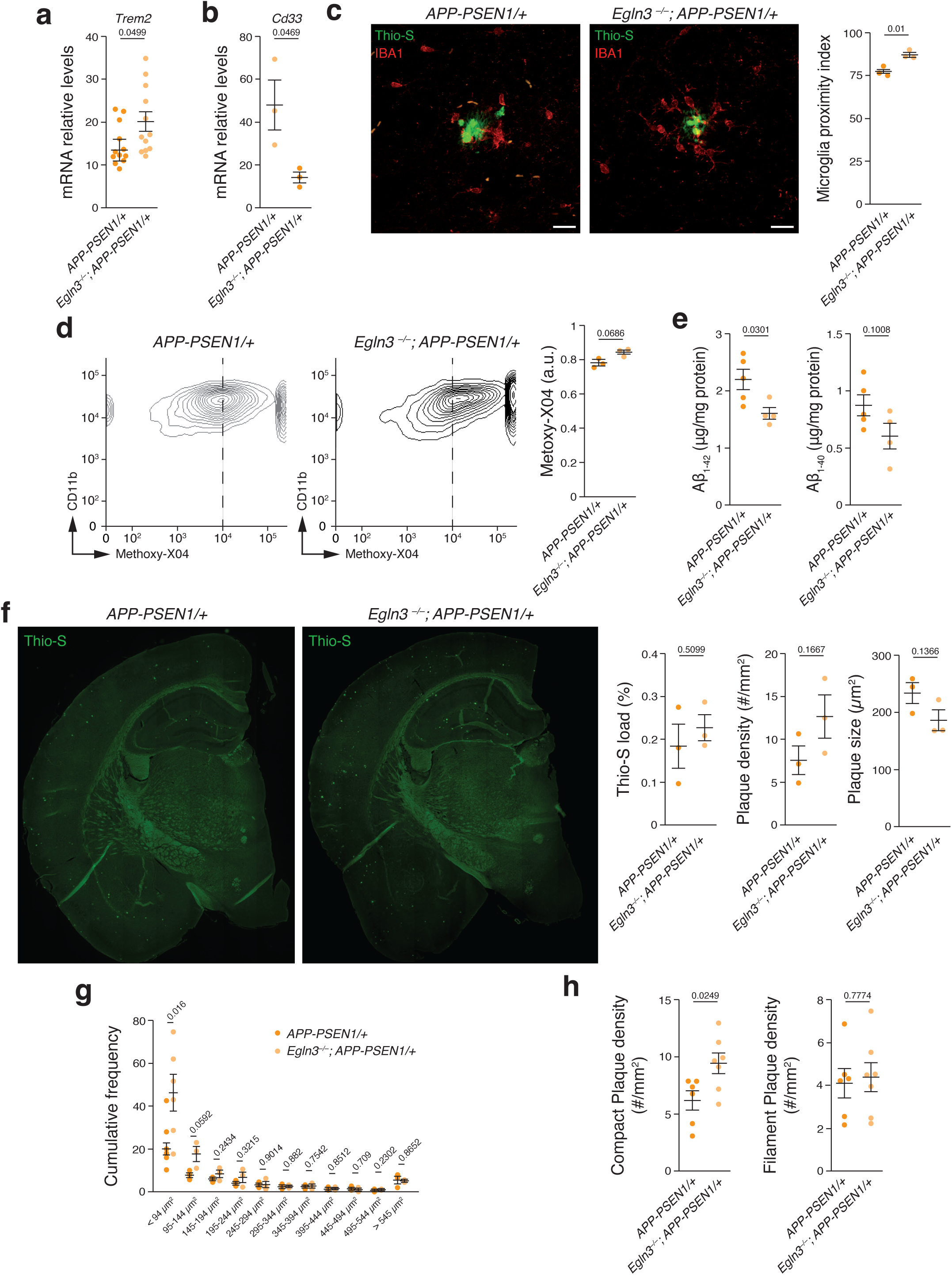
PHD3 limits microglial proximity to Aß plaques. **a**,**b**, *Trem2* and *Cd33* mRNA levels in adult microglia isolated from 12-month-old *APP-PSEN1/+* and *Egln3^−/−^*; *APP-PSEN1/+* mice using qRT-PCR. **c**, Left, representative cortical coronal sections from *APP-PSEN1/+* and *Egln3^−/−^*; *APP-PSEN1/+* mice immunostained with an anti-IBA1 antibody and stained with Thioflavin-S (Thio-S). Scale bars, 20 µm. Right, microglia proximity index: number of IBA1 positive cells in contact with Aß plaques per total microglia within a 40 µm halo around the plaques. **d**, Quantification of Aβ phagocytosis by flow cytometry (left and center panels) of microglia isolated from *APP-PSEN1/+* and *Egln3^−/−^*; *APP-PSEN1/+* 12-month-old mice 3 h after intraperitoneal injection of methoxy-X04. Right panel shows relative measurements. *p* values from two-sided Student’s *t*-test. **e**, Quantification of Aβ_1-42_ and Aβ_1-40_ of half cortical extracts from 6-month-old *APP-PSEN1/+* and *Egln3^−/−^*; *APP-PSEN1/+* mice by ELISA. **f**–**h**, Representative cortical coronal sections from x-month-old *APP-PSEN1/+* and *Egln3^−/−^*; *APP-PSEN1/+* mice stained with Thioflavin-S (Thio-S) (**f**). Right, Thio-S load, plaque density and plaque size (**f**). (**g**) Cumulative frequency of Thio-S plaques. (**h**) Compact and filamentous plaque density (# = number). All data are presented as means ± s.e.m. Individual points in the graphs indicate independent mice used per experiment. *p*-values from two-sided Student’s *t*-test.

As microglial cells act as a physical barrier avoiding the spreading of the axonal pathology caused by Aß^8^, we measured the number of dystrophic neurites found around Aß plaques using a P-TAU antibody, a characteristic marker of this neuronal damage. A substantial decrease in P-TAU labelling was observed around the Aß plaques in the PHD3 mutant mice, indicating an enhanced shielding function of PHD3 deficient microglia (Fig. 6a).

**Figure 6.**
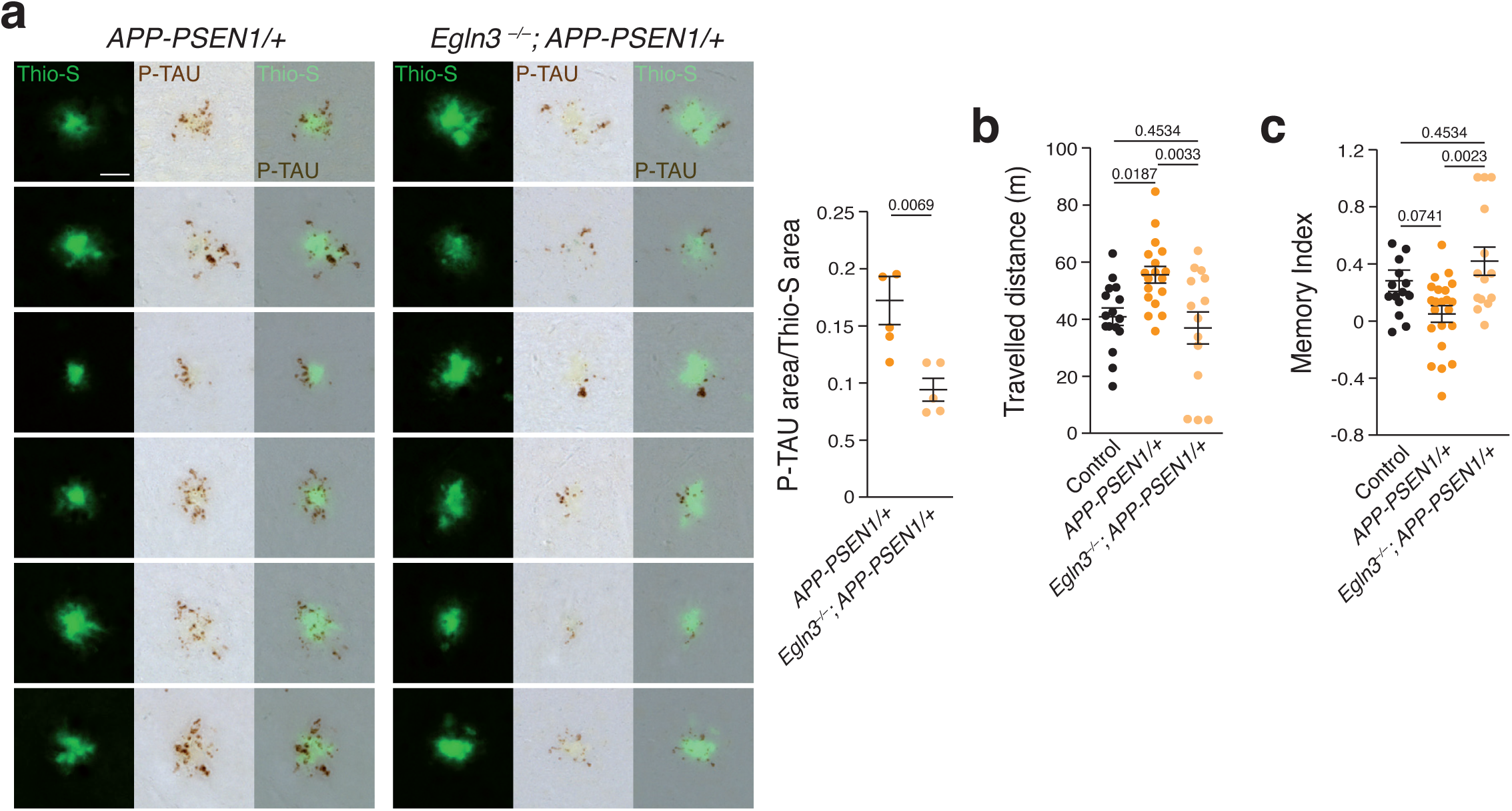
PHD3 deficiency decreases pathology and rescues behavioural deficits of an Aß mouse model. **a**, Left panels, Aß plaques stained with thioflavin-S (Thio-S) and anti-phospho-TAU (P-TAU). Right graph, quantification of the ratio between P-TAU immunoreactive area and Thio-S positive area. *n* = 5 mice (24 plaques per mouse); *p*-values from Student’s *t*-test. **b**,**c**, Spontaneous activity in the open field test (**b**) and memory index (**c**) in the novel object recognition test were measured in 6-month-old mice. (**b**) *n* = 16 control, 18 *APP-PSEN1*, and 14 *Egln3^−/−^*; *APP-PSEN1* mice; (**c**) *n* = 14 control, 21 *APP-PSEN1*, and 14 *Egln3^−/−^*; *APP-PSEN1* mice; *p*-values from ANOVA with Tukey’s post-test. All data are presented as means ± s.e.m. *n* values represent the number of biologically independent experiments.

Last, we investigated whether PHD3 deficiency could improve the cognitive impairment of AD mice. In an open field test, *APP-PSEN1/+* mice exhibited hyperactivity when compared with wild-type and *Egln3^−/−^* control mice (Fig. 6b), as previously described in AD mouse models^10,57^. In sharp contrast, PHD3 deficiency fully rescued this behavioural phenotype (Fig. 6b). To evaluate hippocampal-dependent memory, we used the novel object recognition test. *APP-PSEN1/+* mice showed reduced memory retention when compared with control mice, and the absence of PHD3 fully restored this memory deficit (Fig. 6c). Altogether, our results strongly indicate that PHD3 deficiency improves microglial responses to Aß, decreases local axonal pathology, and recovers behavioural deficits of an AD model.

### Microglial expression of PHD3 is sufficient to induce the IFNS and behavioural alterations

The IFNS is highly relevant for microglial function both during development^46^ or in pathology^20,45^. In both cases, the activation of the IFNS has been linked to signals from neurons that can be detected by microglia, which include dsRNA^46^ and mitochondrial DNA^58^, and downstream pahways^45^. We propose here a new cell autonomous mechanism involved in the activation of the IFNS in microglia, by de-repressing the IFNS through induction of the PHD3-FOXO3 interaction. To prove that the activation of PHD3 in microglia is sufficient to induce the IFNS in the absence of pathology, we developed a new mouse model with overexpression of *Egln3* at the *Rosa26* locus in a CRE-dependent manner (Supplementary Fig. 4a). First, we validated the model by crossing *Rosa26^LSL-Egln3/+^* with a ubiquitous CRE::ERT2 line (*Ubc^Cre::Ert2/+^*), treating mice with tamoxifen, and estimating the degree of *Egln3* overexpression in kidney and brain (Supplementary Fig. 4b). We then crossed the *Rosa26^LSL-Egln3/+^* with the *Cx3cr1^Cre/+^* mouse line (Fig. 7a), which strongly recombines in microglia from development^59^, and verified the upregulation of *Egln3* in isolated microglia (Supplementary Fig. 4c). To investigate if PHD3 expression in the absence of Aß pathology was sufficient to induce the activation of the IFNS, we measured the mRNAs levels of several genes included in the IFNS in mRNA extracted from brain cortex, showing an upregulation of the IFNS (Fig. 7b).

**Figure 7.**
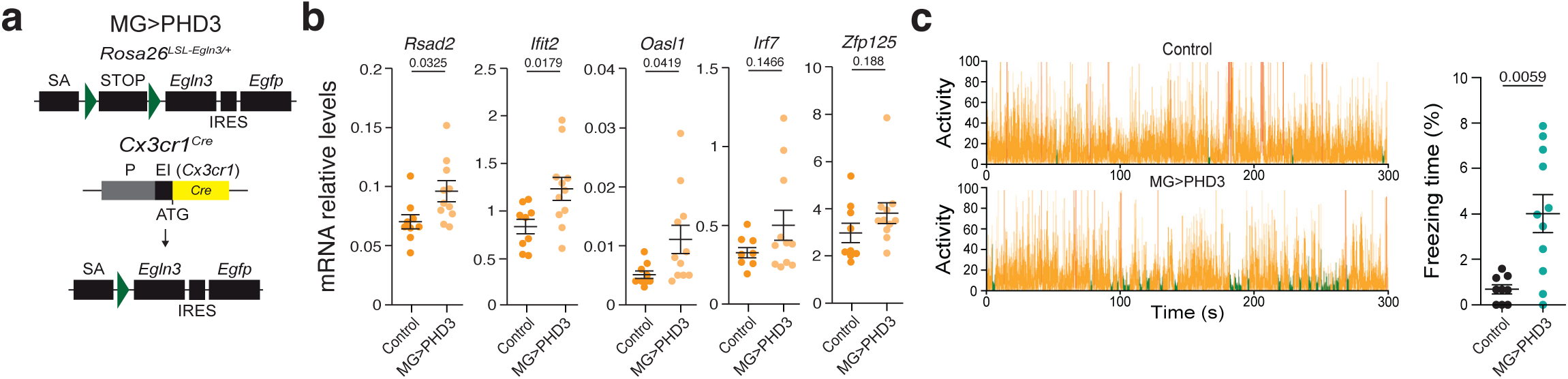
PHD3 overexpression is sufficient to induce the IFNS and behavioural alterations. **a**, Schematic representation (non-scaled) of the *Cx3cr1^CRE/+^*; *Rosa26^LSL-Egln3/+^* (MG>PHD3) mouse model. SA: splicing acceptor. **b**, qRT-PCR analysis of the expression of the IFNS in control and MG>PHD3. **c**, Control and MG>PHD3 mice were recorded in a chamber equipped with a high sensitivity weight detection system for 3 min. Orange, no freezing, and green, freezing. Right graph: freezing time (%). All data are presented as means ± s.e.m. Individual points in the graphs indicate independent 9-month-old mice used per experiment. *p*-values from two-sided Student’s *t*-test.

As microglia has an important role during development^1,2,46^ and we have recently shown that activation of microglia during the perinatal period may associate with behavioural deficits^60^, we measured the freezing time in mice with PHD3 microglial overexpression. We found a clear increase in freezing time when compared to control mice (Fig. 7c), suggesting that PHD3 and its associated IFNS overexpression are sufficient to alter brain function.

## Discussion

We have previously described HIF1 as a master regulator of AD microglial transcription^23^. Here, we show that *Egln3*, encoding for PHD3, is a HIF1 target gene, whose expression is highly restricted to AßAM. *In vitro*, we observe that oAß induces HIF1 and PHD3 expression and correlates with the activation of the IFNS in microglia. We also show that FOXO3 is regulated by PHD3 and a repressor of the IFNS, as previously described in bone marrow macrophages^44^. *In vivo*, PHD3 deficiency leads to a microglial transcriptional reprogramming, characterized by increased DAM and reduced IFNS, which correlated with a better AßAM fitness. PHD3 absence also improves the neuropathology and behavioural deficits of an AD mouse model. Finally, we demonstrated that PHD3 expression in microglia is sufficient to induce the IFNS in the absence of Aß pathology and to cause brain functional alteration, strongly suggesting a cell autonomous mechanism.

We have previously shown that the HIF1-mediated transcriptional program is a characteristic of AßAM^23^, mainly due to the low perfusion of Aß plaques^24^. Although not demonstrated *in vivo*, the role of HIF1 in microglia had been inferred from that in monocyte-derived cells^30,40^, where it has an important contribution in inducing a pro-inflammatory phenotype, characterized by a metabolic switch to glucose fermentation (lactic acid generation) and production of TNF*α* and IL-1ß^28,29^. However, several groups have challenged this presumed HIF1 role in microglia by showing that i) resting microglia is more fermentative than the rest of brain cells^61^ and activation of microglia in physiology^62^ or pathology^23^ associates with upregulation of the mitochondrial electron transport chain encoding genes; ii) hexokinase 2 is almost exclusively expressed by microglia in brain, upregulated in DAM, and its deficiency associates with mitochondrial microglial dysfunction^63^; and iii) genetic overexpression of HIF1 (VHL conditional mutation) in microglia reduces mitochondrial gene expression, DAM, and proliferation^23^. Our data further supports differential consequences for HIF1 accumulation in microglia, with no TNF*α* and IL1ß induction and increased IFNS, unveiling an unexpected role of the hypoxia signalling pathway regulating viral responses in microglia, similar to what was observed in other cell models^64^. However, the two microglial populations express HIF1 and PHD3 in both human^32^ and AD mouse models (^23^ and this report), DAM and IFN microglia, but only the later express the IFNS. Interestingly, although both HIF1 and FOXO3 were proposed as DAM drivers in human iPSCs-derived microglia, simultaneous knockdown of HIF1 and FOXO3 resulted in modestly reduced DAM but elevated IFNS expression^32^, supporting a role for the PHD3-FOXO3 interaction downstream of HIF1, and suggesting that it may gate the transition between these two cellular states. Our data also suggest that transcriptional regulation combined with niche factors (hypoxia and hypoperfusion^24^) contribute to sculp the AßAM responses.

Although no previous studies have linked PHD3 with the activation of anti-microbial responses, PHD3 is elevated in macrophages from patients suffering chronic inflammatory diseases such as Crohn’s disease or ulcerative colitis^33^. Interestingly, the absence of PHD3 has been associated with clinical improvement in mouse models of ulcerative colitis and acute lung injury, through HIF-independent mechanisms^35^. However, it has been shown worse outcome in a HIF1-dependent manner in a sepsis mouse model by increasing the inflammatory responses^34^. Although *Egln3* mRNA is consistently expressed in activated microglia during physiology and pathology, this is the first report showing a role for PHD3 in microglia to our knowledge.

We propose that the IFNS response mediated through the PHD3-FOXO3 pathway in AD microglia is maladaptive, inducing the activation of microglial cells against a non-real threat, such as a viral infection that is not taking place in the brain of AD mouse models. Similar “sterile” (in the absence of pathogen) activation of immune cells has been observed in the peripheral immune system and can be pathogenic^65^. Intriguingly, Aß has recently been shown to have anti-microbial properties^66^, polymorphisms in several IFNS were found associated to AD^67^, other AD risk genes are proposed to activate the IFNS^68^, and IFNS expression is upregulated in AD patients carrying the *TREM2^p.R47H^* variant^69^. The present findings indicate that modulation of the PHD3-FOXO3 pathway in microglia could offer a novel therapeutic target for AD.

## Methods

### Mice

Mice were housed under controlled temperature (22°C) and humidity conditions in a 12 h light/dark cycle with ad libitum access to food and water. Housing and treatments were performed according to the animal care guidelines of the European Community Council (86/60/EEC). All animal procedures were conformed under the Spanish law and approved with numbers 26/04/2016/064; 06/04/2020/050, and 17/10/2023/089 (“Consejería de agricultura, pesca y desarrollo rural. Dirección general de la producción agrícola y ganadera”). B6.Cg-Tg(APPswe,PSEN1Δ9E)85Dbo/J (*APP-PSEN1/+*; stock number 34832)^70^, B6J.B6N(Cg)-Cx3cr1^tm1.1(cre)Jung^/J^71^ (*Cx3cr1^Cre/+^*;, stock number 025524), B6.129-Hif1a^tm3Rsjo^/J (*Hif1a^Flox/Flox^*; stock number 007561)^28^, and B6.Cg-*Ndor1^Tg(UBC-cre/ERT2)1Ejb^*/1J (*Ubq-Cre::ERT2/+*; stock number 007001)^72^ mice were obtained from Jackson Laboratories. *APP_751_SL/+* mice (Sanofi)^73^ were provided by Transgenic Alliance-IFFA-Credo. *Egln3^−/−^* were a generous donation from Prof. Sir Peter J. Racliffe’s laboratory. WT were C57/Bl6. To activate CRE::ERT2-mediated recombination, mice were fed for 30 days with a tamoxifen diet (400 mg tamoxifen citrate/kg; Envigo). All experiments were performed with balanced number of male and female mice. *Rosa26*-LoxP-stop-LoxP-*Egln3* mice were generated using a previously described CRISPR/Cas9 editing method^74^. In the knock-in allele, *Egln3* coding region is linked to an IRES-GFP reporter element and placed downstream of the endogenous Rosa26 promoter connected to its transcript with splice acceptor elements. Conditional expression is achieved by insertion of a loxP-flanked transcriptional stop cassette between the Rosa26 promoter and *Egln3* coding region. *Egln3* murine cDNA (GenBank: AJ310548.1) was subcloned from pCMV-SPORT6 plasmid (Genomics Online #ABIN3831109) in the *BamH*1 and *Xho*I sites of pENTRTM 1A Dual vector (ThermoFisher #A10462) and then inserted by LR reaction in pR26-GFP-Dest plasmid (addgene #74283) using the GatewayTM LR ClonaseTM system (Invitrogen, #11791-019). The resulting plasmid (PHD3-pR26-GFP-Dest) was used as the targeting vector for inserting *Egln3* cDNA into *Rosa26* locus. To achieve Cas9 mediated knock-in into the *Rosa26* locus we used sgRosa26-1 RNA as previously described^74^ from plasmid pU6-sgRosa26-1_CBh-Cas9-T2A-BFP (addgene #64216). Targeting of the construct to the *Rosa26* locus was performed in FVB fertilized zygotes via pronuclear co-injection with a mixture of 50 ng/µl Cas9 protein (PNA BIO Inc #CP01), sgRosa26-1 RNA (10 ng/µl) and PHD3-pR26-GFP-Dest targeting vector (10 ng/µl), in collaboration with the University of Seville animal facility CEA-Oscar Pintado. Founders were screened for cassette insertion with R26F2/SAR^74^ and pR26-fwd/PHD3-rev primer combinations. Correct targeting into *Rosa26* locus was confirmed by PCR amplification of the external 5′ region with r26 fwd/r26rev primers^75^. Cas9 off-target activity was assessed in the founder mice as described previously^74^. Founders were crossed to WT FVB mice to generate F1 animals and inheritance of the knock-in allele was confirmed by PCR amplification and by Sanger sequencing using r26 fwd and r26rev primers.

### Flow cytometry

#### Microglia isolation from adult mouse brains

Mice were anesthetized and transcardially perfused with PBS (Gibco) followed by HBSS (-CaCl_2_/-MgCl_2_) (Gibco) and cortices were dissected and then dissociated using a Tissue Chopper (Vibratome, 800 series). Chemical digestion was performed with a mix of papain (Worthington) (8 U/ml) and DNAse I (Sigma) (80 Kunitz U/ml) followed by a Percoll (GE healthcare) gradient at 90% in PBS (v/v). Cells were stained with primary conjugated monoclonal antibodies CD11b-APC and CD45-PE (eBioscience, 1:200) at 4 °C for 30 minutes. Staining with isotype control-APC and isotype control-PE (eBiosciences, 1:200) was used as a negative control. Both control and experimental samples were simultaneously incubated with anti-CD16/CD32 blocker antibody (eBioscience, 1:200). 7-ADD (BD Pharmigen) was added (1:100) to confirm alive microglial isolation as previously described^8^. Cells were washed and sorted using FACSAria Fusion (Becton Dickinson) flow cytometer and data acquired and analyzed with FACSDiva software 8.0 (Becton Dickinson). The gating strategy and data analysis were performed according to guidelines^76,77^. Debris and dead cells were discarded by forward and side scatter patterns. FSC-A and FSC-H events distribution was used to gate single cells (Supplementary Fig. 2a). Microglia subpopulation selection was performed on a contour density plot scaled to 15% probability and a second microglial subpopulation was considered to be present when more than one independent focus was found with these settings. Percentages are relative to the total number of individual cells.

#### In vivo phagocytosis assay

We calculated the percentage of microglia from total single cells incorporating methoxy-X04 (Tocris Biosience) as previously described^10^. 12-month-ol mice were injected with 10 mg/kg of methoxy-X04 in 50% DMSO/50% NaCl 0.9% intraperitoneally. After 3 h, we isolated microglia using our FACS protocol and cells were analysed in FACSCanto II (Becton Dickinson). Negative threshold for methoxy-X04 was established considering microglia from control mice (not Aß-depositing) and *APP-PSEN1/+* mice without methoxy-X04 injections.

### Cell culture

#### HEK293T cell line cultures

Cells were grown in DMEM medium (Gibco) with 10% fetal bovine serum (Gibco) and 1% penicillin/streptomycin (Gibco) in a water-saturated atmosphere of 5% CO_2_ and 5% air. Cells were detached by trypsinization with 0.25% trypsin-EDTA (Gibco). Cells were always plated at 30-50% confluence to prevent anaerobic conditions.

#### Primary microglial cell cultures

Primary microglial cultures were prepared as previously described^78^ from 1-to 5-days old wildtype or *Egln3^−/−^* mouse brains.

#### In cellulo treatments

DMOG. Cells were incubated for 6 hours or 24 hours with 1 mM (HEK293T cells) or 0.1 mM (primary microglial cells) DMOG (Sigma) dissolved in dimethyl sulfoxide (Sigma). A similar amount of dimethyl sulfoxide was added to control cultures. MG132. Cells were incubated for 4.5 hours or 6 hours with 12,5 μM (HEK293T cells) or 10 μM (primary microglial cell cultures) MG132 (Sigma) dissolved in dimethyl sulfoxide (Sigma). A similar amount of dimethyl sulfoxide was added to control cultures. Small interfering RNAs. Primary isolated microglial cell cultures were transfected with small interfering RNAs (26.67 nM) for 48 hours, using Viromer Blue (Lipocalyx) as a transfection reagent following the manufacturer’s instructions. Transfections. HEK293T cells were transfected with different plasmids at different concentrations for 24 hours, using TurboFect (Thermo Scientific) as a transfection reagent following the manufacturer’s instructions. Plasmids used: pCDNA3 (Thermo Fisher); pCDNA3-PHD1::V5^79^; pCMV6-PHD3::V5^79^; pCMV6-FOXO3::MYC-DDK (Origene); pCMV6-FOXO3(P426A/P437A)::MYC-DDK (this work); pEF1 (Thermo Fisher); and pEF1-HIF1a::DDK (unpublished, generously donated by Prof. Sir Peter J. Racliffe).

### RNA extraction and RT-qPCR

Primary cultures, FACS-isolated microglia and mouse brain and kidney samples. RNA was extracted using TRIzol reagent (Life Technologies). RNA samples were treated with PerfeCTa DNase I (Quanta Biosciences) and copied to complementary DNA (cDNA) using qScript cDNA SuperMix (Quanta Biosciences). cDNA from FACS-isolated microglia was amplified following the protocol described in the section “Microarrays”. Real-Time RT-qPCR was performed for all samples in a ViiA7 Real-Time PCR System (Applied Biosystems) using Power SYBER-Green PCR Master Mix or Taqman (Applied Biosystems) (Supplementary Table 3).

### Protein extraction, immunoprecipitation, and western blot

Primary microglia cell cultures and HEK293T cell lines. For primary microglia cell cultures and brain tissue, total proteins were extracted using TRIzol reagent (Life Technologies) according to the manufacturer’s instructions. An RC-DC protein assay kit (Bio-Rad) was used for quantifications. For HEK293T cells total proteins were extracted using a lysis buffer (100 mM NaCl, 20 mM Tris-HCl pH 7.4, 5 mM MgCl_2_, 0.5% (v/v) Nonidet P-40, 1x proteases inhibitor without EDTA, endonuclease 1:2000). The Pierce 660 Protein Assay Reagent (Thermo Scientific) was used for quantifications. For immunoprecipitation anti-FLAG magnetic beads (Sigma) were used and incubated at 4 °C overnight with the protein extracts. To elute proteins, we used 0.5 M ammonium hydroxide, 0.5 mM EDTA, pH 10. The eluate was neutralized by addition of 10% v/v 1 M Tris pH 7 prior to addition of Laemmli buffer. Western blots were performed using standard procedures. The antibodies used were anti-HIF1*α* (Cayman, 1:500), anti-FOXO3 (Cell Signaling, 1:1,000), anti-V5 Tag (Cell Signaling; 1:1,000), anti-FLAG HRP conjugated (Sigma, 1:2,000) and anti-*β*-actin (Abcam, 1:5,000). HRP-conjugated anti-rabbit (1:10,000) or anti-mouse (1:10,000) antibodies and a Western ECL Substrate kit (Bio-Rad) were used for signal detection.

### *In situ* hybridization and immunostaining

#### Brain In situ hybridization

Brain tissues were cryoprotected in sucrose, embedded in OCT compound (Tissue-Tek) and kept at −80 °C. 20 μm coronal slices were obtained with a cryostat (Leica). The RNAScope 2.5 (ACD) protocol was used to detect *Egln3* mRNA (ACD) according to the manufacturer’s instructions, using a HybEZ oven (ACD). Subsequent immunostaining was performed for microglia staining (with the IBA1 marker) and nuclear staining (DAPI dye). After the RNAScope 2.5 protocol, slices were incubated for 10 minutes in PBS/TRITON X-100 0.3% (v/v) and washed in PBS. Samples were incubated with anti-IBA1 antibody (Wako, 1:500) O/N at 4 °C. Slices were then incubated with Alexa Fluor 488 anti-rabbit (Invitrogen, 1:400) for 1 hour at room temperature and DAPI (Sigma, 1:1000) stained before mounting with Fluoromont-G (Southern Biotech). Images were acquired using a confocal microscope (Leica SP2).

#### Brain immunostaining

The brain was removed from perfused mice with PBS and immediately fixed overnight (15 h) at 4°C with 4% PFA in PBS. The brain was paraffin-embedded using an automatic tissue processor (ASP300S, Leica) and paraffin blocks cut in 20 µm thick coronal sections using a microtome (RM2255, Leica). Immunostaining was performed according to standard protocols. Primary antibodies used: anti-IBA1 (1:500), anti-Aß 6e10 (1:500), anti-P-TAU AT8 (1:500). For immunohistochemistry, Envision+ kit (DAKO) was used for chromatic staining. Secondary antibodies were added the reaction was developed with 3,3-diaminobenzidine (DAB, DAKO). For immunofluorescent studies, we used secondary antibodies anti-mouse or anti-rabbit conjugated with Alexa-488 or Alexa-568. Thioflavin-S, and DAPI staining were used as counterstains.

### Image quantification

#### Microglia density estimation

Unbiased stereological analysis was accomplished by systematic random sampling using a CAST Grid System (Olympus). A 25% of complete hemicortex was sampled with dissectors of 106,954.7 µm^2^ in three slices per mice from –1.7 to –2.06 mm relative to Bregma. Cortical area was manually outlined at 4x magnification. Microglia number in dissectors was quantified by IBA1 marker colocalization with nuclear DAPI staining at 20x magnification. For each hemicortex, total sample area was obtained by multiplying dissector area by number of dissectors. Microglial density was calculated by dividing total microglia number by total sampled area using the Cavaleri method.

#### Microglia proximity index and AßAM number

Proximity index was calculated as the proportion of microglial cells in contact with each Aß plaque considering all microglia closer than 40 µm from Aß plaque’s border. We analyzed randomly selected compact plaques of similar size (15 plaques from three mice per genotype). Sampling area was the whole cortex from –1.94 to –2.3 mm relative to Bregma. Aß plaques was measured by using Fiji software, transforming pictures into 8-bit images, applying a lower threshold of 0 and an upper threshold of 255, and using analyze particle tool. The region of interest generated after amyloid plaques measurement by Fiji was submitted to enlarge function in order to extend its periphery 40 µm. All microglia whose soma was within the extended halo or touching its periphery were quantified by IBA1 marker colocalization with nuclear DAPI staining. Separately, microglia directly in contact with the Aß plaque and within this halo were also quantified. The ratio between microglia directly in contact with Aß plaque and total microglia number in each extended halo constituted the microglial proximity index.

#### Microglial, Thio-S, and Aß load

Brain slices were stained with either IBA1, Thio-S, or an anti-Aß antibody. Measurements were performed in superimages, blind to the genotypes. Three cortex superimages per mouse were obtained at 10x magnification from –1.7 to –2.3 mm relative to Bregma with New CAST BX61 (Olympus) and using the same microscope setting for all the samples. Fiji software was used to analyze the superimages by transforming them into 8-bit, selecting the cortical area of interest manually, and subtracting areas with evident noise or artifacts like folded tissue. We created a segmented binary mask by using the same lower and upper thresholds all along the samples for each staining. Analyze particles function was used to calculate the total area covered by all the particles contained in the segmented image. The load was defined as the percentage of occupancy by the signal divided the total cortical area studied and multiplied by 100. We also estimated the number of plaques per area (density) and the averaged plaque area.

#### Microglia morphology

Individual microglial cells from the dorso-lateral cortex (12 cells from three slices per mouse) were randomly pictured using the same acquisition parameters. Images consisted in 18 Z-stacks in an Olympus BX61 microscope at 100x magnification, from –1.70 and –2.06 mm relative to Bregma. Maximum intensity projections were obtained for further analysis. We counted the number of primary branches (emerging directly from the soma), and the number of intersections (Scholl index) with a circumference of 667.91 µm^2^. We also calculated the percentage of area covered by individual microglial cells in circumference of 1,108.24 µm^2^ with and without the soma (a circumference of 155.02 µm^2^).

Amyloid plaques were visualized with anti-Aß and Thio-S staining and were randomly selected blind to the treatment in the cortex. Quantifications were done in superimages generated with the NewCAST system (Visiopharm) associated with the microscope BX61 (Olympus). The number of microglia cells surrounding amyloid plaques was determined after immunostaining for IBA1 and staining with DAPI using immunofluorescence. Microglia coverage of individual amyloid plaques was obtained by normalizing IBA1 occupied area by the area occupied for the corresponding Thio-S reactive plaque, calculated from binary masks generated with appropriate thresholds for all images in Fiji. Results are presented as a percentage of IBA1 per Aß plaque area. In wild-type and regions distal to Aß plaques, full images were quantified and a density of the marker was calculated. *Microglial stereology*. The measurements were performed blind to the treatment. Unbiased stereological approach using an Olympus BX61 microscope combined with the CAST system. The sample area was then manually outlined and the total area quantified using CAST software and microglia between two specific bregma points were estimated using a dissector area of 28,521.3 μm2 (CAST). The dentate gyrus was chosen as a sample area.

#### Compact and filamentous Aß plaque density

All Thio-S positive plaques bigger than 55 µm2 were quantified in the whole cortex and classified as filamentous, if no dense core was visible, or compact plaques, in which a solid and compact core was identifiable. Plaques were visualized a 20 x magnification in CAST Grid System (Olympus).

#### P-TAU load

Plaques were randomly selected by Thio-S staining and the Thio-S and P-TAU area were estimated using Fiji in individual plaques. Results are presented as a percentage of P-TAU area per amyloid plaque area.

### Microarray

RNA quality was analyzed using an Agilent 2100 Bioanalyzer (Agilent). Only samples with RNA integrity number higher than 7 were used. The total RNA extracted from FACS-isolated microglia was amplified and labeled using GeneChip WT Pico Reagent Kit (Thermo Scientific). The amplified cDNA was quantified, fragmented and labeled for hybridization to GeneChip Mouse Transcriptome 1.0 Array (Thermo Scientific) using 5.5 μg of single-stranded cDNA product and following protocols outlined in the user manual. Washing, staining (GeneChip Fluidics Station 450, Thermo Scientific) and scanning (GeneChip Scanner 3000, Thermo Scientific) were performed following the protocols outlined in the user manual. Raw data from the Expression Console extraction software (Thermo Scientific) were imported into the statistical program R (RStudio Inc.) using the LIMMA/Bioconductor package^80^. The quality of the data was assessed using Array Quality Metrics package^81^. Data were normalized using Robust Multi-Array method and differential expression analysis was performed using the LIMMA/Bioconductor package^82^. Gene expression data from *APP-PSEN1/+*, *5xfAD*, *APP_751_SL/+*, and *Egln3^−/−^* mouse models were analyzed by GSEA using Biological Processes C5-v5.2, KEEG and the custom IFNS and FOXO3_BS^44^.

### Human snRNAseq data

Dataset. The single-cell RNA sequencing data used in this study was sourced from the publicly available dataset hosted at https://compbio.mit.edu/microglia_states/from^32^. The dataset, comprising both the metadata and gene expression count matrix, was downloaded in RDS format. To facilitate downstream analysis in Python, the RDS files were converted into CSV format for metadata and MTX format for the count matrix, enabling compatibility with standard Python-based computational tools for single-cell data analysis. Data preprocessing. Quality control metrics were applied to remove outlier cells from the dataset. Cells were filtered based on the total number of counts, retaining those with at least 600 and no more than 9,000 counts. Additionally, cells were required to express a minimum of 500 genes and a maximum of 3,000 genes. To exclude cells with excessive mitochondrial content, a mitochondrial fraction cutoff of 0.1% was applied. The initial dataset contained 174,420 cells, and following quality control, 168,373 cells remained. Raw count data was normalized using the normalize_per_cell() and log1p() functions from the Scanpy library^83^ [2] to standardize gene expression levels for subsequent analyses. Clustering analysis. To replicate the cluster definitions from the original study^32^, we used the Seurat clusters provided in the metadata, as the original cluster labels were not available. These Seurat-defined clusters were correlated with the highly expressed gene markers for each microglial state, as reported in the Supplementary Table of the original article. While most clusters could be mapped to the original labels, some were challenging in terms of clear correlation. For clusters that could not be confidently matched, we classified them as Non-Defined (ND). Consequently, we defined specific microglial states, including microglia (MG)0, MG2, MG3, MG4, MG5/barrier-associated macrophages (BAM), MG6, MG7, MG8, MG10, MG11, MG12 and T-cells. IFNS and FOXO3_binding_sites (BS) signatures analysis. A predefined list of IFN and FOXO3 gene signatures from mice was applied to assess the correlation between these signatures and the microglial states in our single-cell dataset. For each cell, we calculated an expression score for both the IFN and FOXO3_BS signatures, based on the mean expression level of all genes associated with each signature. These calculated values, referred to as IFNS and FOXO3-GS, were then analyzed using the rank_genes_groups() function from Scanpy^83^, which applies the Wilcoxon rank-sum test. This approach allowed us to determine whether any of the signatures could serve as significant markers for any of the microglial states.

### Behavioral analysis

#### Locomotion activity

Mice were filmed in an open field arena of 45 x 45 cm using an automatic tracking software (SMART). Mice were recorded for 15 min following the protocol from the International mouse phenotyping resource of standardized screens (IMPRESS; https://www.mousephenotype.org/impress/ProcedureInfo?procID=81). Travelled distance was calculated using the SMART software 3.0 (Panlab-Harvard Apparatus).

#### Novel object recognition test

Short-term memory was evaluated following a previously published protocol^84^. Two objects were presented to each mouse for 15 min in a 45 x 45 cm arena. After one hour, the mouse was exposed to one of the previous objects being the other replaced by a novel one. A short memory index was calculated by subtracting the number of approaches to the novel object minus the number of approaches to the old object and divided by the total number of approaches. Recording and analyzing software was SMART 3.0.

#### Freezing behavior

The duration of freezing was estimated using a fully automatic startle and fear conditioning system (Panlab). Mice were introduced in a chamber equipped with a high sensitivity weight transducer system and the immobility was measured using the PACWINCSFR software (Panlab), with a lower bound threshold of 18% and a minimal duration of 1 s.

### Aß ELISA

Frozen hemicortices were homogenized with a Dounce grinder (Sigma) in 4x w/v solution of PBS containing protease inhibitor cocktail (Roche; 1:25) and phosphatase inhibitor (Roche; 1:100). Samples were subjected o consecutive centrifugation steps to separate soluble from insoluble fractions. First, samples were centrifuged at 600 g for 5 min to pellet the nuclear fraction. Next, the supernatants were ultracentrifuged at 100,000 g for 1 h at 4°C to isolate PBS-insoluble (pellets) from PBS-soluble (supernatants) proteins. The insoluble fraction was resuspended in 8.2 M guanidine-HCl, 50 mM Tris-HCl, diluted to 5 M guanidine-HCl, and incubated at room temperature for 4 h with occasional inversion to promote dissolution. Total protein concentration was quantified with RC-DC kit (Bio-Rad) and Aß_1-40_ and Aß_1-42_ concentrations were tested using human Aß40 and Aß42 ELISA kits (Invitrogen).

### Statistical analysis

All individual measurements constitute independent biological replicates. Samples with *n ≥* 9 were evaluated for normal distribution using D’Agostino and Pearson’s omnibus normality test. Samples with *n* < 9 were analyzed using parametric tests. Comparisons between two groups were performed with two-sided unpaired Student’s t-test, whereas comparisons between more than two groups were done with analysis of variance (ANOVA) with Tukey’s test. Data are expressed as means ± s.e.m. *p ≤* 0.05 was considered statistically significant. Statistical analyses and graphs were performed in GaphPad Prism (GraphPad).

---

**Supplementary Figure 1. The AßAM from mouse models express the IFN microglial response**. **a,b**, Gene set enrichment analysis (GSEA) in adult microglia isolated from *5xfAD/+*, *APP-PSEN1/+*, and *APP_751_SL/+* (**a** and **b**) Aß-depositing mouse models. Left panels, enrichment plots; Right panels show the heatmap of the top 30 ranking leading edge genes. **c**, HEK-293T cells transfected with both PHD1-V5 or PHD3-V5 and an empty (EV) or HIF1*α*-FLAG (HIF1*α*) vector; and incubated with DMOG for 24 h. Upper panels show western blot with anti-FLAG (FOXO3) and bottom panels with anti-V5 (PHD1 and PHD3) with proteins extract (INPUT) and after immunoprecipitation (IP) with an anti-FLAG antibody. **d**, HEK-293T cells transfected with both PHD3-V5 or an empty (EV), and a FOXO3-FLAG vector; and incubated with vehicle (C), DMOG (D) or MG132 (MG) for 24 h. Western blots with anti-FLAG (FOXO3) and anti-V5 (PHD3) in proteins extracts (INPUT) and after immunoprecipitation (IP) with an anti-FLAG antibody.

**Supplementary Figure 2. PHD3 deficiency abolishes the anti-viral transcriptional response of AßAM. a**, Debris and dead cells were discarded by forward (FSC) and side (SSC) scatters dispersion of events (far left panel). Singlets of events were selected according to FSC height (FSC-H) versus area (FSC-A; left panel). 7-ADD staining was performed as control to test alive microglial cells isolation (right panel). Microglial cells, positive for CD45 and CD11b markers, were selected. **b,c**, GSEA of adult microglia from 12-month-old *Egln3^−/−^*; *APP-PSEN1/+ versus APP-PSEN1/+* (left panels, enrichment plots). Right panels show the heatmap of the top-ranking leading-edge genes. Red symbolizes upregulation and blue, downregulation.

**Supplementary Figure 3. Histological characterization of the PHD3 deficient mouse model**. **a**, Number (#) of microglial cells surrounding Aß plaques in *APP-PSEN1/+* and *Egln3^−/−^*; *APP-PSEN1/+* mice. **b**, Representative cortical coronal sections from *APP-PSEN1/+* and *Egln3^−/−^*; *APP-PSEN1/+* mice stained with IBA1. Right, microglial (IBA1) load. **c**, Morphologic characterization of non-plaque associated microglia from *APP-PSEN1/+* and *Egln3^−/−^*; *APP-PSEN1/+* mice and stained with IBA1. **d**, Representative cortical coronal sections from *APP-PSEN1/+* and *Egln3^−/−^*; *APP-PSEN1/+* mice immunostained with an anti-Aß antibody. Right, Aß load.

All data are presented as means ± s.e.m. Individual points in the graphs indicate independent 12-month-old mice used per experiment. *p*-values from two-sided Student’s *t*-test.

**Supplementary Figure 4. Characterization of the *Rosa26^LSL-Egln3/+^* mouse model**. **a**, Schematic representation (non-scaled) of the *Rosa26^LSL-Egln3^* mouse model generation. SA: splicing acceptor. LHA: left homology arm. RHA: right homology arm. **b**, qRT-PCR analysis of the expression of the *Egln3* mRNA levels in *Ub^CRE::ERT2/+^*; *Rosa26^LSL-Egln3/+^* (UB>PHD3)mouse model. **c**, qRT-PCR analysis of the expression of the *Egln3* mRNA levels in microglia (CD11b^+^) and other cells (CD11b^−^) isolated from control and MG>PHD3 mouse model.

All data are presented as means ± s.e.m. Individual points in the graphs indicate independent mice used per experiment. *p*-values from two-sided Student’s *t*-test.

## ACKNOWLEDGMENTS

We thank Prof. Sir Peter J. Ratcliffe for hosting N.L.-U. in his laboratory for PHD-FOXO3 interaction experiments. We also wish to thank K. Levitsky (microscopy), M.J. Castro (flow cytometry), F.J. Moron (genomics), and R. Duran (histology) for advice and technical assistance in experiments at IBiS core facilities. R.M.-D. was the recipient of a “Sara Borrell” fellowship from ISCIII (CD09/0007), M.A.S.-G., N.L.-U., C.O-dSL., C. R.-M., and S.Q-C. were the recipient of an FPU fellowship from Spanish Ministry of Education, Culture, and Sport (respectively, FPU13/00530, FPU14/02115, AP2010-1598, FPU16/02050, and FPU22/00974), A.H.-G. was the recipient of an FPI fellowship from Spanish Ministry of Education, Culture, and Sport (BES-2010-033886) and B.M.-R. was the recipient of a “Junta de Andalucía” predoctoral fellowship (2021). A.E.R.-N. was recipient of JdlC-F fellowship from MCIN/AEI/ 10.13039/501100011033 (FJCI-2015-23708). Work was supported by grants to A.P./J.V./A.M.M.-C. by MCIN/AEI/ 10.13039/501100011033, ISCIII (FORT23/00008 –A.P.– and PI21/00915 –J.V.–), and FEDER (RTI2018-096629-B-100, PID2021-126894OB-I00 and “FEDER Una manera de hacer Europa”, SAF2017-90794-REDT, and PIE13/0004), by the regional Government of Andalusia (“Proyectos de Excelencia” P12-CTS-2138, P20_01312; Biotechnology Plan applied to Health PRTR-C17.I1; FEDER Andalucia 2014-2020, US-1261055) co-funded by CEC, REC_EU, and FEDER funds, and by the “Ayuda de Biomedicina 2018”, Fundación Domingo Martínez.

## Extended and Supplementary Data files

Extended Data Figs. 1 to 4

Supplementary Data Tables 1 to 3

## Data availability

Original data is provided as Source Data files in the Supplementary Data, indicating the correspondence with each main and extended figure. Transcriptomics data are available from the Gene Expression Omnibus Dataset

## Author contributions

M.S.-G., N.L.-U., R.M.-D., A.B.M.-M., A.E.R.-N., and A.P. conceived and designed the research. M.S.-G., N.L.-U., R.M.-D., C.O-d.S.L., J.M.B-R., S.Q.-C., D.C.-R., A.M.M.-C., B.M.-R., C.R.-M., A.H-G., V.N., A.E.R.-N., and A.P. performed the experiments. M.S.-G., N.L.-U., R.M.-D., C.O-d.S.L., J.M.B-R., A.M.M.-C., M.V., J.V., M.C., A.E.R.-N., and A.P. analysed the data. J.L.-B. and M.C. contributed mouse models/samples/techniques. M.A.S.-G., A.E.R.N., and A.P. wrote the manuscript.

## References

1. Tay, T. L., Savage, J. C., Hui, C. W., Bisht, K. & Tremblay, M. È. Microglia across the lifespan: from origin to function in brain development, plasticity and cognition. J. Physiol. 595, 1929–1945 (2017).

2. Prinz, M., Jung, S. & Priller, J. Microglia Biology: One Century of Evolving Concepts. Cell 179, 292–311 (2019).

3. Pimenova, A. A., Raj, T. & Goate, A. M. Untangling Genetic Risk for Alzheimer’s Disease. Biol. Psychiatry 83, 300–310 (2018).

4. Serrano-Pozo, A., Gómez-Isla, T., Growdon, J. H., Frosch, M. P. & Hyman, B. T. A Phenotypic Change But Not Proliferation Underlies Glial Responses in Alzheimer Disease. Am. J. Pathol. 182, 2332–2344 (2013).

5. Condello, C., Yuan, P., Schain, A. & Grutzendler, J. Microglia constitute a barrier that prevents neurotoxic protofibrillar Aβ42 hotspots around plaques. Nat. Commun. 6, 6176 (2015).

6. Leyns, C. E. G. et al. TREM2 function impedes tau seeding in neuritic plaques. Nat. Neurosci. 22, 1217–1222 (2019).

7. Wang, Y. et al. TREM2-mediated early microglial response limits diffusion and toxicity of amyloid plaques. J. Exp. Med. 213, 667–675 (2016).

8. Yuan, P. et al. TREM2 Haplodeficiency in Mice and Humans Impairs the Microglia Barrier Function Leading to Decreased Amyloid Compaction and Severe Axonal Dystrophy. Neuron 90, 724–739 (2016).

9. Heneka, M. T., Golenbock, D. T. & Latz, E. Innate immunity in Alzheimer’s disease. Nat. Immunol. 16, 229–36 (2015).

10. Heneka, M. T. et al. NLRP3 is activated in Alzheimer’s disease and contributes to pathology in APP/PS1 mice. Nature 493, 674–8 (2013).

11. Lucin, K. M. & Wyss-Coray, T. Immune Activation in Brain Aging and Neurodegeneration: Too Much or Too Little? Neuron 64, 110–122 (2009).

12. Sarlus, H. & Heneka, M. T. Microglia in Alzheimer’s disease. J. Clin. Invest. 127, 3240– 3249 (2017).

13. Hickman, S. E. et al. The microglial sensome revealed by direct RNA sequencing. Nat. Neurosci. 16, 1896–905 (2013).

14. Krasemann, S. et al. The TREM2-APOE Pathway Drives the Transcriptional Phenotype of Dysfunctional Microglia in Neurodegenerative Diseases. Immunity 47, 566–581.e9 (2017).

15. Keren-Shaul, H. et al. A Unique Microglia Type Associated with Restricting Development of Alzheimer’s Disease. Cell 169, 1–15 (2017).

16. Orre, M. et al. Isolation of glia from Alzheimer’s mice reveals inflammation and dysfunction. Neurobiol. Aging 35, 2746–60 (2014).

17. Holtman, I. R. et al. Induction of a common microglia gene expression signature by aging and neurodegenerative conditions: a co-expression meta-analysis. Acta Neuropathol. Commun. 3, 31 (2015).

18. Friedman, B. A. et al. Diverse Brain Myeloid Expression Profiles Reveal Distinct Microglial Activation States and Aspects of Alzheimer’s Disease Not Evident in Mouse Models. Cell Rep. 22, 832–847 (2018).

19. Taylor, J. M. et al. Type-1 interferon signaling mediates neuro-inflammatory events in models of Alzheimer’s disease. Neurobiol. Aging 35, 1012–1023 (2014).

20. Roy, E. R. et al. Type I interferon response drives neuroinflammation and synapse loss in Alzheimer disease. J. Clin. Invest. 130, 1912–1930 (2020).

21. Sala Frigerio, C., et al. The Major Risk Factors for Alzheimer’s Disease: Age, Sex, and Genes Modulate the Microglia Response to Aβ Plaques. Cell Rep. 27, 1293–1306.e6 (2019).

22. Olah, M. et al. Single cell RNA sequencing of human microglia uncovers a subset associated with Alzheimer’s disease. Nat. Commun. 11, (2020).

23. March-Diaz, R. et al. Hypoxia compromises the mitochondrial metabolism of Alzheimer’s disease microglia via HIF1. *Nat*. aging 1, 385–399 (2021).

24. Alvarez-Vergara, M. I. et al. Non-productive angiogenesis disassembles Aß plaque-associated blood vessels. Nat. Commun. 12, 3098 (2021).

25. Morita, M. et al. MTORC1 controls mitochondrial activity and biogenesis through 4E-BP-dependent translational regulation. Cell Metab. 18, 698–711 (2013).

26. Ulland, T. K. et al. TREM2 Maintains Microglial Metabolic Fitness in Alzheimer’s Disease. Cell 170, 649–663.e13 (2017).

27. Kaelin, W. G. & Ratcliffe, P. J. Oxygen Sensing by Metazoans: The Central Role of the HIF Hydroxylase Pathway. Mol. Cell 30, 393–402 (2008).

28. Cramer, T. et al. HIF-1α is essential for myeloid cell-mediated inflammation. Cell 112, 645– 657 (2003).

29. Tannahill, G. M. et al. Succinate is an inflammatory signal that induces IL-1β through HIF-1α. Nature 496, 238–242 (2013).

30. Baik, S. H. et al. A Breakdown in Metabolic Reprogramming Causes Microglia Dysfunction in Alzheimer’s Disease. Cell Metab. 30, 493–507.e6 (2019).

31. Wendeln, A.-C. et al. Innate immune memory in the brain shapes neurological disease hallmarks. Nature 556, 332–338 (2018).

32. Sun, N. et al. Human microglial state dynamics in Alzheimer’s disease progression. Cell 186, 4386–4403.e29 (2023).

33. Escribese, M. M. et al. The Prolyl Hydroxylase PHD3 Identifies Proinflammatory Macrophages and Its Expression Is Regulated by Activin A. J. Immunol. 189, 1946–1954 (2012).

34. Kiss, J. et al. Loss of the oxygen sensor PHD3 enhances the innate immune response to abdominal sepsis. J. Immunol. 189, 1955–65 (2012).

35. Walmsley, S. R. et al. Prolyl hydroxylase 3 (PHD3) is essential for hypoxic regulation of neutrophilic inflammation in humans and mice. J. Clin. Invest. 121, 1053–63 (2011).

36. Swain, L. et al. Prolyl-4-hydroxylase domain 3 (PHD3) is a critical terminator for cell survival of macrophages under stress conditions. J. Leukoc. Biol. 96, 365–375 (2014).

37. Beneke, A. et al. Loss of PHD3 in myeloid cells dampens the inflammatory response and fibrosis after hind-limb ischemia. Cell Death Dis. 8, e2976–e2976 (2017).

38. Zheng, X. et al. Prolyl hydroxylation by EglN2 destabilizes FOXO3a by blocking its interaction with the USP9x deubiquitinase. Genes Dev. 28, 1429–1444 (2014).

39. Rodriguez, J. et al. Substrate-Trapped Interactors of PHD3 and FIH Cluster in Distinct Signaling Pathways. Cell Rep. 14, 2745–2760 (2016).

40. Peruzzotti-Jametti, L. et al. Mitochondrial complex I activity in microglia sustains neuroinflammation. Nature 628, 195–203 (2024).

41. Metzen, E. et al. Intracellular localisation of human HIF-1α hydroxylases:implications for oxygen sensing. J. Cell Sci. 116, 1319–1326 (2003).

42. Ginouvès, A., Ilc, K., Macías, N., Pouysségur, J. & Berra, E. PHDs overactivation during chronic hypoxia ‘desensitizes’ HIFalpha and protects cells from necrosis. Proc. Natl. Acad. Sci. U. S. A. 105, 4745–4750 (2008).

43. Wang, Y. et al. TREM2 lipid sensing sustains the microglial response in an Alzheimer’s disease model. Cell 160, 1061–71 (2015).

44. Litvak, V. et al. A FOXO3–IRF7 gene regulatory circuit limits inflammatory sequelae of antiviral responses. Nature 490, 421–425 (2012).

45. Minter, M. R. et al. Deletion of the type-1 interferon receptor in APPSWE/PS1ΔE9 mice preserves cognitive function and alters glial phenotype. Acta Neuropathol. Commun. 4, 72 (2016).

46. Escoubas, C. C. et al. Type-I-interferon-responsive microglia shape cortical development and behavior. Cell 187, 1936–1954.e24 (2024).

47. Honda, K. et al. IRF-7 is the master regulator of type-I interferon-dependent immune responses. Nature 434, 772–777 (2005).

48. Cockman, M. E. et al. Lack of activity of recombinant HIF prolyl hydroxylases (PHDs) on reported non-HIF substrates. Elife 8, 1–27 (2019).

49. Saunders, A. et al. Molecular Diversity and Specializations among the Cells of the Adult Mouse Brain. Cell 174, 1015–1030.e16 (2018).

50. Bishop, T. et al. Abnormal sympathoadrenal development and systemic hypotension in PHD3-/-mice. Mol. Cell. Biol. 28, 3386–400 (2008).

51. Rangaraju, S. et al. Differential phagocytic properties of CD45low microglia and CD45high brain mononuclear phagocytes-activation and age-related effects. Front. Immunol. 9, (2018).

52. Crotti, A. & Ransohoff, R. M. Microglial Physiology and Pathophysiology: Insights from Genome-wide Transcriptional Profiling. Immunity 44, 505–15 (2016).

53. Guerreiro, R. et al. TREM2 Variants in Alzheimer’s Disease. N. Engl. J. Med. 368, 117– 127 (2013).

54. Jonsson, T. et al. Variant of TREM2 associated with the risk of Alzheimer’s disease. N. Engl. J. Med. 368, 107–16 (2013).

55. Griciuc, A. et al. Alzheimer’s Disease Risk Gene CD33 Inhibits Microglial Uptake of Amyloid Beta. Neuron 78, 631–643 (2013).

56. Colonna, M. & Wang, Y. TREM2 variants: new keys to decipher Alzheimer disease pathogenesis. Nat. Rev. Neurosci. 17, 201–207 (2016).

57. Walker, J. M. et al. Spatial learning and memory impairment and increased locomotion in a transgenic amyloid precursor protein mouse model of Alzheimer’s disease. Behav. Brain Res. 222, 169–175 (2011).

58. Di Domizio, J. et al. Nucleic acid-containing amyloid fibrils potently induce type I interferon and stimulate systemic autoimmunity. Proc. Natl. Acad. Sci. 109, 14550–14555 (2012).

59. Goldmann, T. et al. A new type of microglia gene targeting shows TAK1 to be pivotal in CNS autoimmune inflammation. Nat. Neurosci. 16, 1618–1626 (2013).

60. Mora-Romero, B. et al. Microglia mitochondrial complex I deficiency during development induces glial dysfunction and early lethality. Nat. Metab. (2024). doi:10.1038/s42255-024-01081-0

61. Bernier, L. P. et al. Microglial metabolic flexibility supports immune surveillance of the brain parenchyma. Nat. Commun. 11, (2020).

62. He, D. et al. Disruption of the IL-33-ST2-AKT signaling axis impairs neurodevelopment by inhibiting microglial metabolic adaptation and phagocytic function. Immunity 55, 159–173.e9 (2022).

63. Hu, Y. et al. Dual roles of hexokinase 2 in shaping microglial function by gating glycolytic flux and mitochondrial activity. Nat. Metab. 4, 1756–1774 (2022).

64. Hwang, I. I. L., Watson, I. R., Der, S. D. & Ohh, M. Loss of VHL confers hypoxia-inducible factor (HIF)-dependent resistance to vesicular stomatitis virus: role of HIF in antiviral response. J. Virol. 80, 10712–23 (2006).

65. Rock, K. L., Latz, E., Ontiveros, F. & Kono, H. The sterile inflammatory response. Annu. Rev. Immunol. 28, 321–42 (2010).

66. Kumar, D. K. V. et al. Amyloid-β peptide protects against microbial infection in mouse and worm models of Alzheimer’s disease. Sci. Transl. Med. 8, 340ra72 (2016).

67. Salih, D. A. et al. Genetic variability in response to amyloid beta deposition influences Alzheimer’s disease risk. Brain Commun. 1, 1–13 (2019).

68. Ramdhani, S. et al. Tensor decomposition of stimulated monocyte and macrophage gene expression profiles identifies neurodegenerative disease-specific trans-eQTLs. PLOS Genet. 16, e1008549 (2020).

69. Korvatska, O. et al. Triggering Receptor Expressed on Myeloid Cell 2 R47H Exacerbates Immune Response in Alzheimer’s Disease Brain. Front. Immunol. 11, 1–16 (2020).

70. Jankowsky, J. L. et al. Mutant presenilins specifically elevate the levels of the 42 residue beta-amyloid peptide in vivo: evidence for augmentation of a 42-specific gamma secretase. Hum. Mol. Genet. 13, 159–70 (2004).

71. Yona, S. et al. Fate Mapping Reveals Origins and Dynamics of Monocytes and Tissue Macrophages under Homeostasis. Immunity 38, 79–91 (2013).

72. Ruzankina, Y. et al. Deletion of the Developmentally Essential Gene ATR in Adult Mice Leads to Age-Related Phenotypes and Stem Cell Loss. Cell Stem Cell 1, 113–126 (2007).

73. Blanchard, V. et al. Time sequence of maturation of dystrophic neurites associated with Aβ deposits in APP/PS1 transgenic mice. Exp. Neurol. 184, 247–263 (2003).

74. Chu, V. T. et al. Efficient generation of Rosa26 knock-in mice using CRISPR/Cas9 in C57BL/6 zygotes. BMC Biotechnol. 16, 4 (2016).

75. Nyabi, O. et al. Efficient mouse transgenesis using Gateway-compatible ROSA26 locus targeting vectors and F1 hybrid ES cells. Nucleic Acids Res. 37, (2009).

76. Herzenberg, L. a, Tung, J., Moore, W. a, Herzenberg, L. a & Parks, D. R. Interpreting flow cytometry data: a guide for the perplexed. Nat. Immunol. 7, 681–5 (2006).

77. Orre, M. et al. Acute isolation and transcriptome characterization of cortical astrocytes and microglia from young and aged mice. NBA 35, 1–14 (2014).

78. Sanchez-Mejias, E. et al. Soluble phospho-tau from Alzheimer’s disease hippocampus drives microglial degeneration. Acta Neuropathol. 132, 897–916 (2016).

79. Masson, N. et al. The HIF prolyl hydroxylase PHD3 is a potential substrate of the TRiC chaperonin. FEBS Lett. 570, 166–170 (2004).

80. Ritchie, M. E. et al. limma powers differential expression analyses for RNA-sequencing and microarray studies. Nucleic Acids Res. 43, e47–e47 (2015).

81. Kauffmann, A., Gentleman, R. & Huber, W. arrayQualityMetrics--a bioconductor package for quality assessment of microarray data. Bioinformatics 25, 415–416 (2009).

82. Phipson, B., Lee, S., Majewski, I. J., Alexander, W. S. & Smyth, G. K. Robust hyperparameter estimation protects against hypervariable genes and improves power to detect differential expression. Ann. Appl. Stat. 10, 946–963 (2016).

83. Wolf, F. A., Angerer, P. & Theis, F. J. SCANPY: large-scale single-cell gene expression data analysis. Genome Biol. 19, 15 (2018).

84. Leger, M. et al. Object recognition test in mice. Nat. Protoc. 8, 2531–7 (2013).

