## Supplementary figures S1-S4 for "The PHD3-FOXO3 axis modulates the interferon type I response in microglia aggravating Alzheimer’s disease progression"

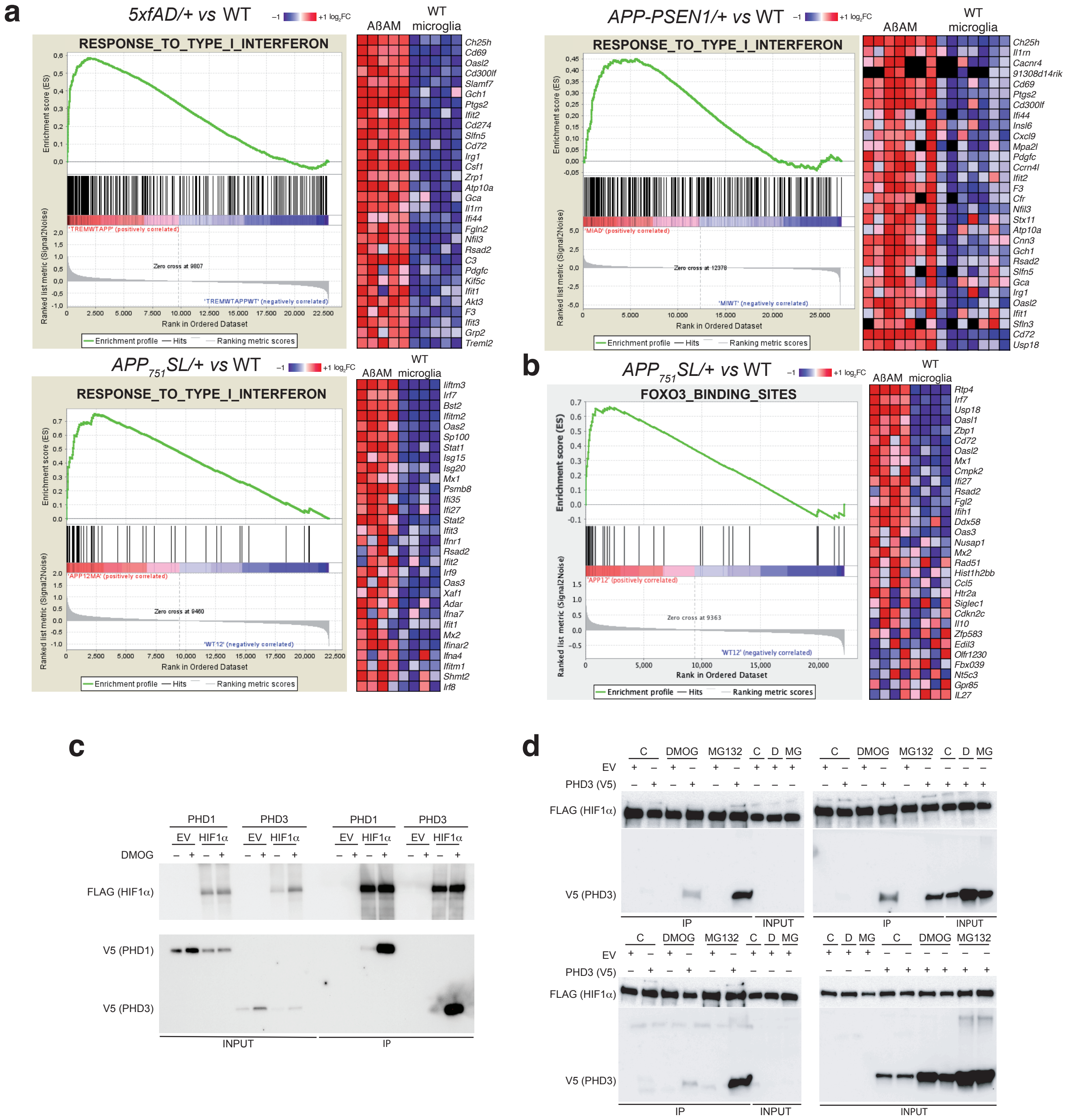


Sanchez-Garcia *et al.*, Supplementary Figure 1

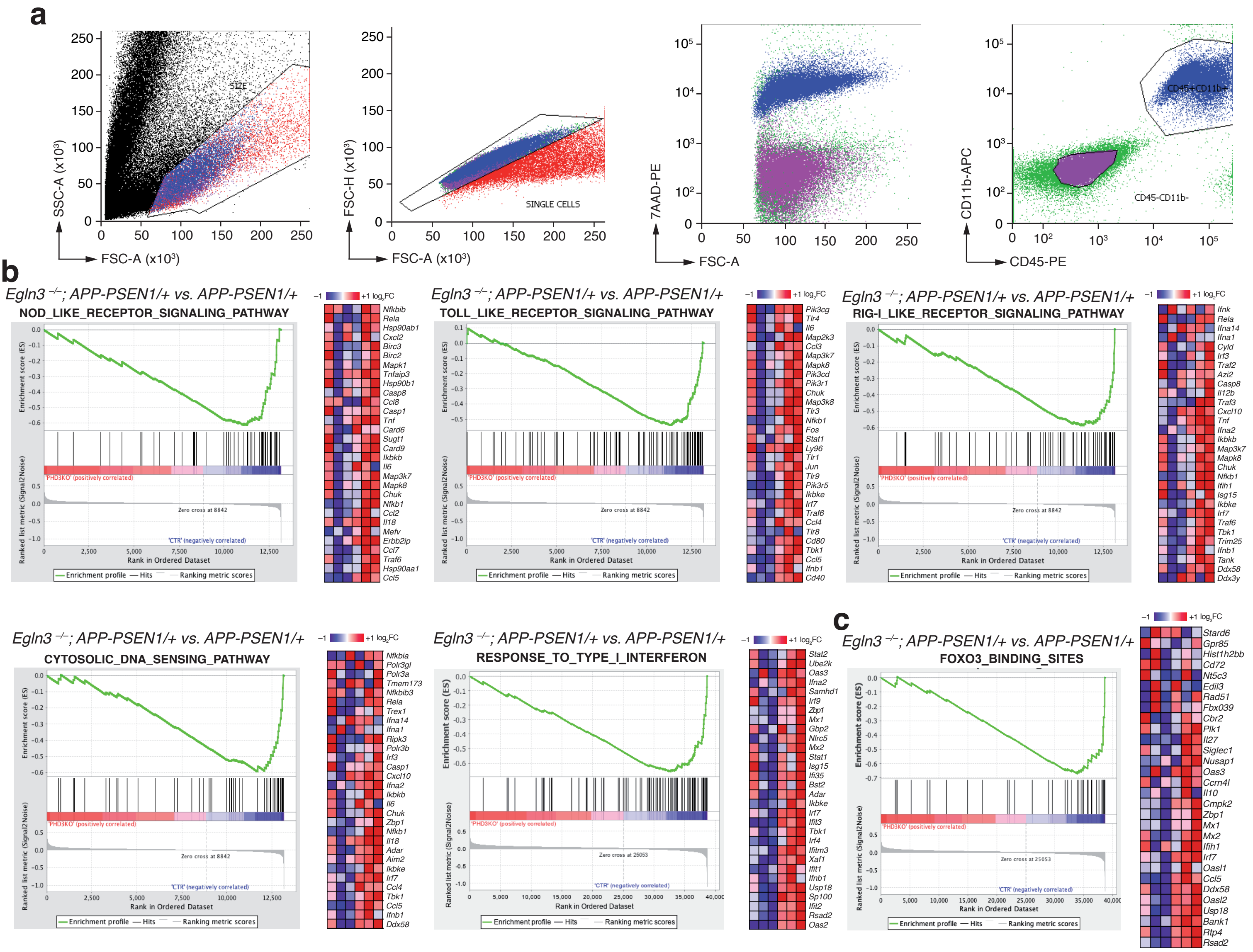


Sanchez-Garcia *et al.*, Supplementary Figure 2

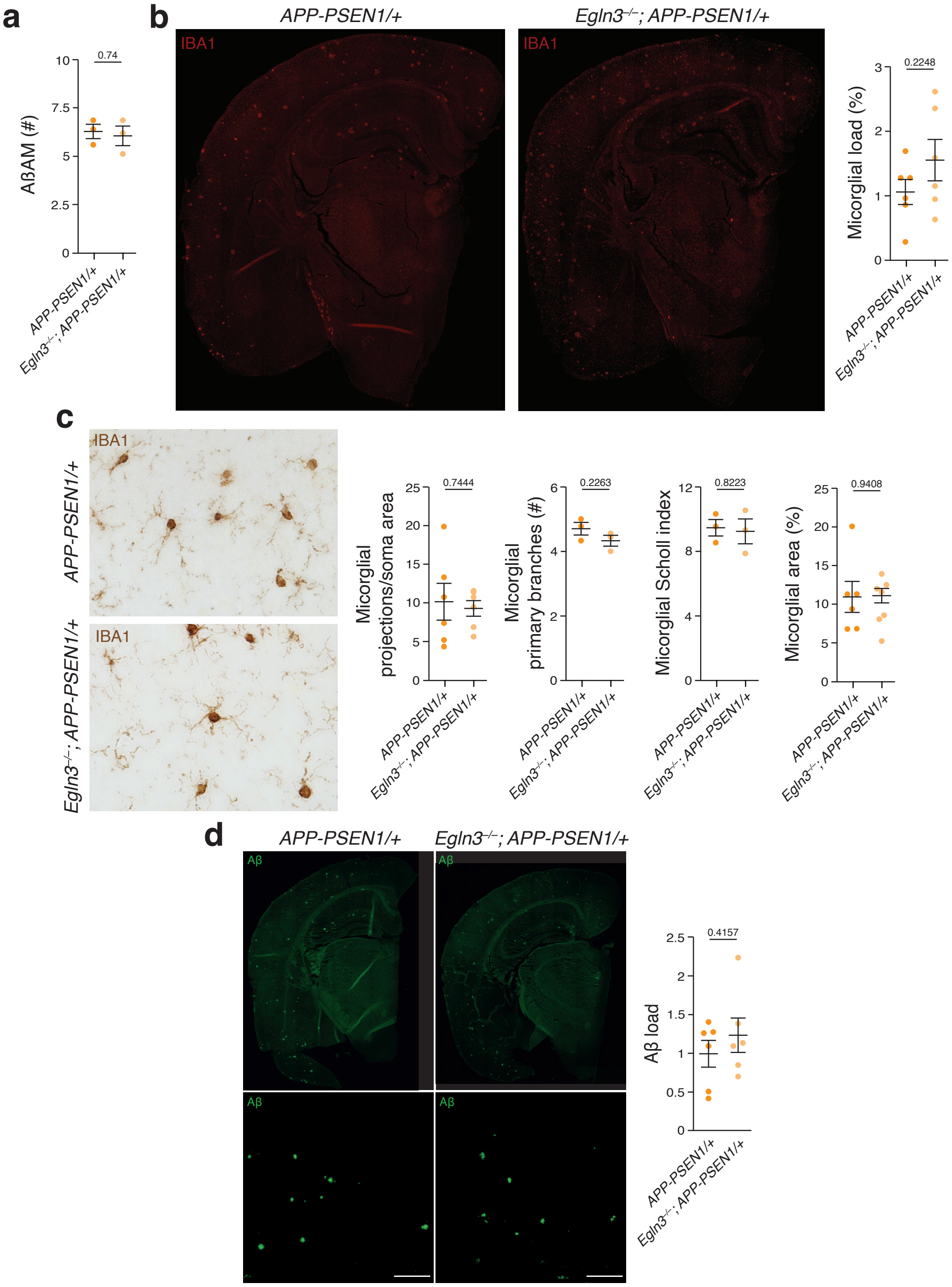


Sanchez-Garcia *et al.*, Supplementary Figure 3

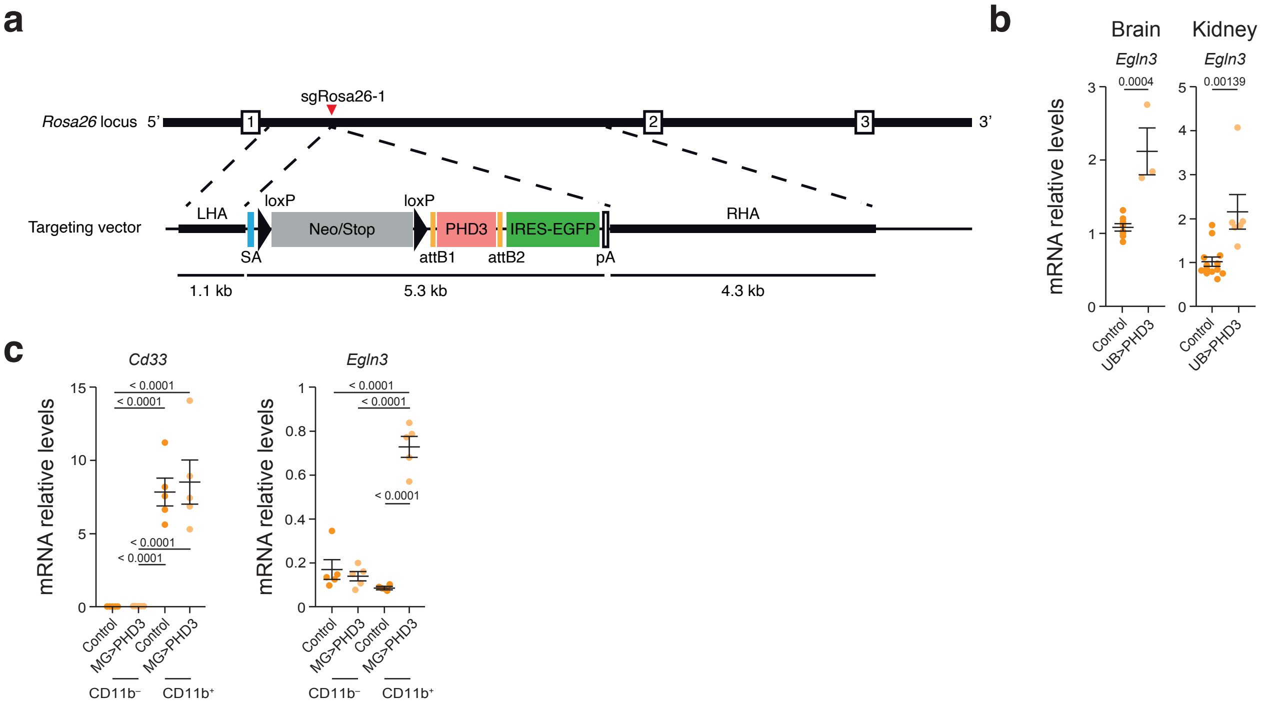


Sanchez-Garcia *et al.*, Supplementary Figure 4
